# Zooplankton feeding behavioral signatures in the morphology of macroscale prey spatial distribution

**DOI:** 10.1101/2025.08.27.672564

**Authors:** Eduardo H. Colombo, Corina E. Tarnita, Juan A. Bonachela

## Abstract

The problem of pattern and scale remains a central problem in ecology, bridging fundamental and applied questions. Marine microbial communities are a case in point. For instance, to understand the role of zooplankton in oceanic biogeochemistry, their response to changes in environmental conditions, and the implications for ecosystem services (e.g., fisheries), it is critical to understand zooplankton trophic interactions and how they change in a rapidly changing climate. This understanding, however, remains elusive because, unlike for phytoplankton, for which remote sensing of macroscale patterns can provide insight into their microscale dynamics and community composition, obtaining this information for zooplankton largely rests on quantifying the difficult-to-monitor microscale interactions among millions of individuals with different behaviors, and between individuals and their environment. Here, we investigate whether it is possible to obtain indirect information on zooplankton from the macroscale spatial distribution of their prey. To tackle this “problem of scale”, we develop a rigorous coarse-graining methodology that connects individual-level properties with macroscale spatial patterns. We demonstrate that the shape of the prey spatial distribution can indeed encode information about zooplankton feeding behavior and community dynamics. Specifically, we predict a change in dominant feeding behavior—from non-motile to motile feeding—as one moves from areas of high to areas of low prey density. These results thus suggest a novel application for remote sensing approaches: the potential tracking of consumer behavioral signatures in the large-scale patterns of the resource. Importantly, the scaling-up methodology that we developed to check whether those signatures exist is general, and can be used to link scales rigorously and systematically in any system in which the complexity of individual dynamics makes connecting scales intractable.

---

Marine ecosystems stand on the shoulders of microbial organisms. Phytoplankton—microbial primary producers—have a protagonist role as they recycle atmospheric carbon dioxide to produce more than half the oxygen we breath every day, sequester carbon to the deep ocean in the form of sinking marine snow, and introduce carbon and other essential nutrients (e.g., nitrogen, phosphorus) into the marine food web [1–3]. Zooplankton, predators of phytoplankton and other prey (e.g., bacteria, other zooplankton), and preferential resource for fish, create a link that connects the lower to the higher trophic levels of the marine food web, as well as to the deep sea via excretion of dissolved and particulate organic matter [4, 5]. Similarly to their prey, zooplankton are part of diverse communities with different functional roles, and thus have different contributions to ecosystem dynamics. Understanding and predicting oceanic biogeochemistry and other key marine ecosystem services, thus, requires understanding and predicting what species dominate the different levels of the microbial food web.

A tool available to determine the presence and even composition of large phytoplankton communities is remote sensing which, using chlorophyll as a proxy, has revealed mesmerizing kilometer-scale phytoplankton spatial patterns (see Fig. 1) with impressive spatial and temporal resolution [6, 7]. Remote sensing, however, cannot in general directly provide information about other key players in the microbial food web (see [8] for a rare exception). Standard approaches to obtaining information about zooplankton and other important actors of oceanic carbon flow, such as viruses, involve *in-situ* observations—a much greater effort compared to remote sensing as it entails ship expeditions, large interdisciplinary collaborations, and substantial financial resources per case studied (e.g., [9]). Despite the large amount of data they can produce, these available methods can only offer limited information about species interactions or zooplankton feeding behavior [10]. Measuring and characterizing feeding behavior *in situ*, for example, is challenging without affecting (and thus biasing) observations [11]. Behavior, however, influences feeding rates, therefore affecting important ecological outcomes such as the flux of carbon towards the deep ocean and the structure of the microbial marine food web [12, 13]. Thus, these limitations ultimately handicap the understanding of, and ability to predict, oceanic biogeochemical processes and how they change across space and time [14, 15].

**Figure 1:**
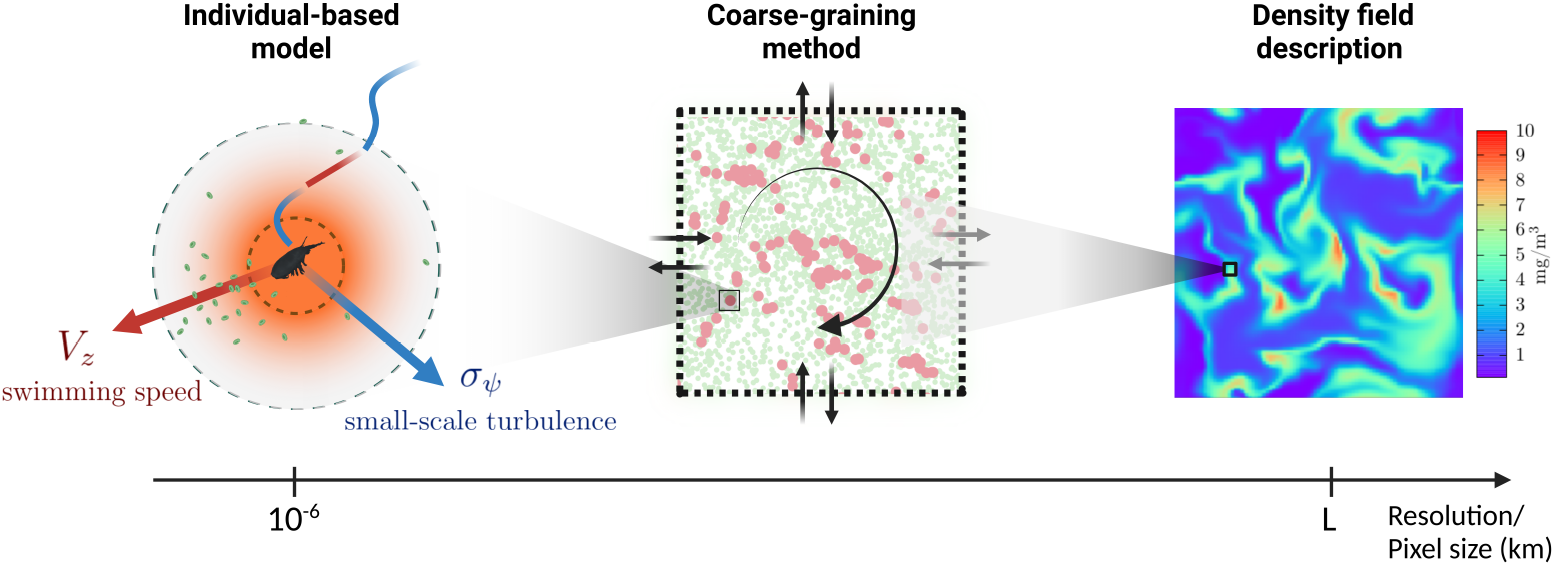
From the micro-to macroscale. Using our individual-based model as a starting point (left panel), we performed a coarse-graining method (middle panel) to obtain the interaction and transport terms for a density-field description (right panel). Our methodology provides a connection across scales of observation that facilitates understanding the large-scale consequences of the individual-level features such as the swimming speed of the focal zooplankton species, *V*_*z*_, and the typical speed stemming from small-scale turbulence, *σ*_*ψ*_ (see *Materials and methods*).

One way to address these limitations is to search for complementary indirect information in remote-sensing data [16–18]. Extracting information on zooplankton dynamics and feeding behavior in a non-invasive way using satellite images of, for example, macroscopic (kilometer-scale) patterns of phytoplankton or other prey would be an important first step to further our understanding of their role in those patterns. This is, however, a non-trivial and potentially infeasible endeavor due to the complexity of the links between the observed large-scale patterns and the individual-level interactions that influence them [19].

If this ‘problem of scale’ is generally difficult to solve for any given biological system, it is especially challenging in the case of zooplankton because of their feeding behavior. Although, for decades, macroscopic models used the simplifying assumption that zooplankton are passive relative to oceanic turbulence [20, 21], advances in high-resolution computer simulations [22] and lab-scale experiments [12, 23, 24] have stressed that zooplankton do not just ‘go with the flow’[25]: a wide variety of zooplankton can sense the surrounding environment and use this information to decide where to swim [26]. As a result, a tug-of-war emerges between this impressive behavioral capability and turbulence, influencing where and when prey are eaten. At the microscale, this tug-of-war could lead to the formation of zooplankton patches [23, 25, 27] that potentially also affect the spatial distribution of e.g., dissolved organic matter (resulting from zooplankton excretion) or particulate organic matter (from zooplankton sloppy feeding), all of which could influence and contribute to the emergent large-scale patterns as well as have rippling effects across the rest of the marine food web [13]. However, the mechanistic underpinnings of these patterns—i.e. whether, how, and what individual-level behaviors are responsible for shaping the patterns at different scales—are still unknown. This is in large part due to the technical intractability of connecting, step-by-step, the individual-level zooplankton behavior (well-known thanks to controlled experiments, e.g. [28]) with the associated microscale formations and, ultimately, with the resulting macroscale prey patterns, connection that involves a broad spatial range (from millimeters to kilometers) and massive numbers of individuals to be tracked.

Despite this missing connection, existing theories have been able to reproduce qualitatively some features of large-scale oceanic plankton patterns by resorting to generic arguments or heuristic approximations that did not require explicitly linking individual and macroscopic levels [20, 21, 29–31]. For example, the fact that zooplankton and phytoplankton form distinct patterns has been captured by simple models with local facilitation and long-range competition among planktonic populations (e.g. [20]). Similarly, other studies have explored the role of zooplankton in the carbon flux (e.g., assessed consumption rates) by using phenomenological population-level expressions that depend on individual traits [17, 28, 32, 33]. In all previous theories, however, the lack of an explicit mechanistic connection between individual-, micro-, and macroscale properties leaves a causal gap that prevents the utilization of macroscale patterns to infer key aspects of the microbial community, such as the zooplankton feeding behavior that dominates. This causal gap also precludes theories able to predict potential shifts in the community over time, especially as anthropogenic effects promote substantial changes in ocean properties (e.g., toxicity [34], increased temperature, lowered viscosity, stratification [35] and turbidity [36, 37]) that can result in behavioral changes at the individual level.

Here, we attempt to close this causal gap by introducing a novel computational coarse-graining method that allowed us to scale up an individual-based model to a macroscale density-field description (see Fig. 1a). We show that zooplankton feeding behavior leaves fingerprints in prey patterns at multiple scales. At the microscale, the individual-level model reveals that zooplankton active search for resources (active behavior, i.e. swimming towards perceived prey) induces an expected increase in spatial correlation between prey and zooplankton, but also an unexpected correlation among zooplankton, which ultimately form clumps. At large scales, our results show that the shape of the prey patterns can reflect the underlying feeding behavior, and thus that remote sensing can indeed be used to infer behavior: passive (i.e. non-swimming) zooplankton lead to smooth prey patterns while active zooplankton produce sharp pattern edges. We further extended our theoretical model to allow for the two types of behavior to explicitly compete for prey, which uncovered a change of dominance, from passive to active feeding, as the density of prey declines. Such information could be used to, for example, identify the dominant zooplankton species (or, at least, the dominant behavior) across time and environmental gradients, and monitor shifts that might occur under climate change and that would ultimately influence the dynamics of the microbial community [11, 38] and the rest of the marine food web.

## I. RESULTS AND DISCUSSION

We developed a general modeling framework that rigorously connects microscale dynamics to emergent macroscale patterns. The description of the macroscale arises from a coarse-graining method that considers the individual-based dynamics occurring in square domains (pixels) of size *L* that are stitched together, while keeping pixel size below the typical Kolmogorov scale of the ocean [39]. For scales below the pixel size, the individual-level dynamics are described by an individual-level model (IBM) where oceanic turbulence is effectively accounted for by considering that the velocity of both zooplankton and prey individuals is random with standard deviation set by parameter *σ*_*ψ*_ (proxy for turbulence intensity at these scales). For scales beyond *L*, the coarse-graining method uses the IBM to derive a partial differential equation for predator-prey dynamics, using a point-vortex model to describe turbulence with intensity *ψ* (see *Materials and methods* and Fig. S1). Below, we discuss first the microscale dynamics and subsequently the coarse-graining procedure and emergent macroscale properties. Henceforth, we use zooplankton and predator indistinctly.

### A. Microscale dynamics

The IBM accounts for the main processes generally observed across zooplankton and prey types (individual birth and death, zooplankton feeding behavior) and an external flow that aims to mimic the effects of turbulence. Feeding determines prey death rate and zooplankton reproduction rate, and is influenced by the interplay between physics (i.e. flow) and ecology (i.e. predator swimming behavior). At the scale of the IBM, the turbulence flow is represented by a random velocity field that moves individuals; this flow tends to break correlations between prey and zooplankton. Zooplankton can perceive visual and physico-chemical cues (e.g., hydrodynamic disturbances and chemical trails), to which they respond in different ways [28]. In our model, we generically captured feeding behavior by defining a perceptual range *R*_*p*_ and two types of responses to cues within that range: active and passive. Active zooplankton individuals sense prey individuals within the perception range, swim towards them at a speed *V*_*z*_ > 0 (if speed is strong enough for the predator to overcome the flow), and have a probability to catch and ingest the target [40] when the prey is within a catching range *R*_*c*_ < *R*_*p*_. Passive zooplankton individuals, on the other hand, do not swim (i.e. *V*_*z*_ = 0) even if they might sense prey, but they ingest prey if the flow brings both prey and predator within catching range. Active predators in our model would thus resemble cruise feeders that intentionally swim towards prey, while passive predators would encompass zooplankton feeding modes that rely on currents to bring prey within catching range (e.g. ambush feeders) [28]). For further details, see IBM description in Fig. 1a, *Materials and methods*, and *Supplementary Materials*.

Although overall zooplankton behavior depends on various parameters (see above), here we focused on the interplay between flow and swimming behavior. To this end, we varied the small-scale turbulence intensity, *σ*_*ψ*_, and the swimming speed *V*_*z*_. Note that, although swimming entails a metabolic cost, we did not include it in this version of the model for the sake of simplicity. We did so, however, in an extension of the model specifically devised to contrast costs and benefits of feeding behavior (see adaptive-feeding model in *Materials and methods*).

#### Emergent timescale separation

Transport and demographic processes can both introduce spatial correlations and thus play a role in the emergent spatial pattern: the former, via physical spatial rearrangements of individuals; the latter, via changes in species densities (i.e. due to mortality/feeding, which decreases locally the number of individuals, or due to replication, which increases it). The model revealed, however, that transport and demography lead to correlations at different timescales. Given that typical speeds for flow and our generic zooplankton were on the order of ~1cm/s and that the predator perceptual range is ~1cm (see *Materials and methods*), transport-induced spatial features emerged in our simulations in just a few seconds. This timescale was much shorter (approximately 1/10) than the typical time interval between demographic events in the model. As shown in Fig. S2, such a timescale separation occurs for a wide range of prey densities. Importantly, this emergent timescale separation between the short-term relocation of individuals and demographic events allowed us to study these processes independently, as explained in the next two sections.

#### On the short timescale, the movement of active zooplankton induces microscale patchiness

To understand the role of a given swimming speed *V*_*z*_ in the formation of microscale patchiness we focused on the short timescales, where transport occurs in the absence of demographic processes. Thus, we assumed a constant density of prey and zooplankton (*p* and *z*, respectively), initially randomly distributed across space, and we explored a range of values for *V*_*z*_, leaving the other behavioral traits (e.g., perception range) constant. Fig. 2a illustrates the contrast between the typical patterns that emerged, after a short transient, for passive (*V*_*z*_ *=* 0 cm/s) and active (*V*_*z*_ > 0) zooplankton for low values of prey density. For the passive case, the turbulence-controlled transport maintained a random spatial distribution for both species. For active zooplankton, however, the directional movement towards prey increased the correlation between the location of zooplankton (red dots) and prey (green dots) and, unexpectedly, also among zooplankton individuals (for details, see species pair correlation function in Fig. S3). The latter correlation resulted from predators that were sufficiently close to each other and therefore partially shared the same cues, which led them to move in the same direction.

**Figure 2:**
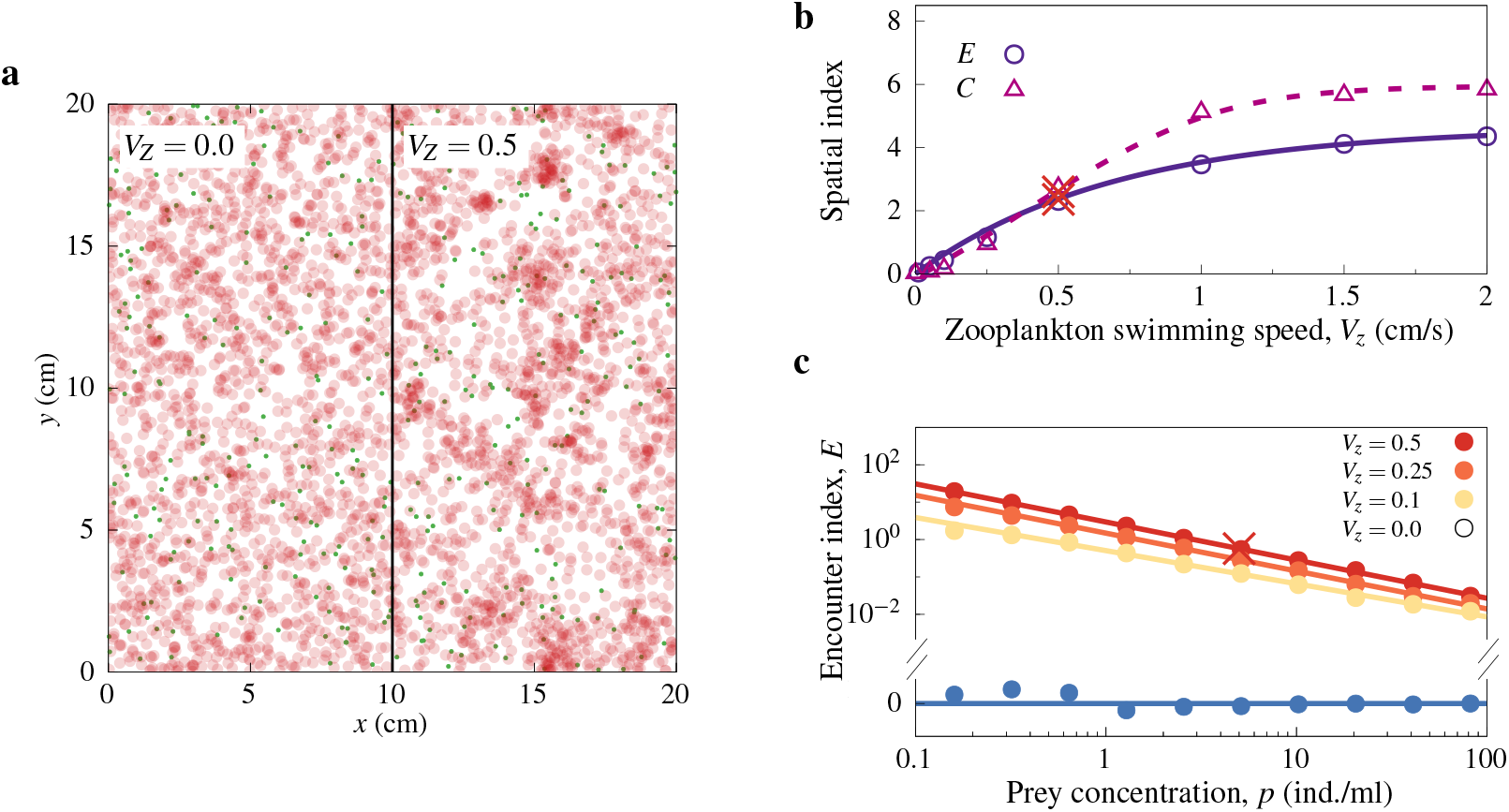
Microscale patchiness emerging from the individual-level model. (a) Spatial distribution for prey (green dots) and predators (80%-transparent red dots) for densities (*p, z*) = (1.28, 5); note that this example was chosen to illustrate the emergence of patterns, which are more pronounced at low prey densities. The left half of the picture shows the passive zooplankton case (*V*_*z*_ = 0.0 cm/s), and the right half the active zooplankton case (*V*_*z*_ = 0.5 cm/s), for easy visual comparison. (b) The patterns from the scenario in panel (a) are characterized by the encounter (*E*) and clumping (*C*) indices, shown here for different zooplankton swimming speeds, *V*_*z*_. (c) Encounter index as a function of prey density for different swimming speeds *V*_*z*_. In both (b) and (c), dots represent the mean across replicates of values reached in the stationary regime, with crosses corresponding to the cases depicted in panel (a). Solid lines are given by Eq. 1. All cases were obtained for moderate turbulence (*σ*_*ψ*_ = 0.25 cm/s).

To quantify the formation of microscale patchiness, we defined two spatial indices: the encounter index, *E* (*t*), related to prey-zooplankton encounters, and the clumping index, *C* (*t*), indicative of zooplankton aggregation (see *Materials and methods*). We defined both indices to be zero in the well-mixed (random) scenario and exhibit positive values in the presence of microscale patchiness (Eq. 6). Fig. S4a-b illustrates how these spatial indices started at a zero value and, after an initial short transient (5 − 10s), reached (quasi-)stationary values, *E* and *C*, which are our values of interest henceforth.

These stationary values, which characterize the microscale patchiness for the given densities (*p, z*), depended on predator swimming speed (Fig. 2b) and turbulence (Fig. S4c-f). For high values of *σ*_*ψ*_, at any fixed *V*_*z*_, turbulence generated displacements that were large relative to the zooplankton catching range, *R*_*c*_, which resulted in random mixing (*E* ~ 0, *C* ~ 0). At moderate values of *σ*_*ψ*_, *E* and *C* increased with *V*_*z*_ and eventually reached a plateau when swimming offset turbulence, as predators could reach and stay close to selected prey targets (Fig. 2b). At low turbulence, i.e. displacements smaller than the catching range, *E* and *C* increased with the swimming speed, but showed a maximum at a finite speed 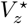 (Fig. S4c-d). For the remainder of our results, we focus on realistic speeds, which are below 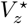 (see *Materials and methods*; we also explored what underlies this interesting peak via a simple model, see *Supplementary Materials* for details). Altogether, these results show that the tug-of-war between mixing (caused by the turbulent flow) and aggregation (caused by feeding behavior) determines species encounters (Fig. S4e,f) and, consequently, potentially influences population-level interactions.

#### Microscale patchiness is density-dependent

Because they result from the relocation of individuals, the patterns emerging from the short-timescale processes above influence species interactions and the dynamics of species densities occurring at the longer timescale. In turn, demographic changes locally add and remove individuals and therefore affect short-term spatial correlations, how many individuals are transported, and the directions that active zooplankton will move into on the shorter timescale. This feedback loop shapes the emergent microscale patterns and how they change over time.

To investigate whether and how prey and predator demography influences the dynamics of microscale patchiness, we considered a given turbulence-speed scenario, i.e. fixed (*σ*_*ψ*_, *V*_*z*_), and monitored (*p*(*t*), *z*(*t*)) over time. Due to the timescale separation, individuals quickly rearranged in space between demographic events, which allowed us to define a stationary encounter and clumping index for each density, i.e. *E* = *E*(*p, z*) and *C* = *C*(*p, z*). We found that, in fact, both indices were significantly correlated only with prey density, and not with zooplankton density, i.e. *E* = *E*(*p*) and *C* = *C*(*p*) (Figs. S5).

Fig. 2c shows the emergent correlation between the encounter index and prey density under different swimming speeds. High prey densities led to frequent encounters with zooplankton similarly to the random encounters expected in the well-mixed scenario (i.e. *E →* 0 as *p → ∞*); as prey density decreased, however, encounters became rare unless the predator actively swam towards prey, and therefore active feeding substantially increased encounters relative to the passive case. Active feeding also increased the fraction of time predators had prey within reach (see *Supplementary Materials*). Both the encounter and the clumping indices (Fig. S5) show that active feeding promotes aggregation at low prey densities, but does not provide any significant advantage at high densities.

From Fig. 2c we infer that the encounter index scales as a power-law with prey density and depends on both small-scale turbulence and zooplankton swimming speed, i.e.:

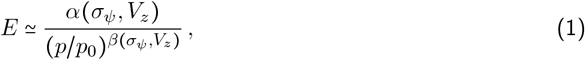

where *p*_0_ = 1 ind./cm^2^ is a (fixed) scaling factor (see Fig. 2c) and the coefficients *α* and *β*, both of which depend on turbulence and swimming speed (*σ*_*ψ*_, *V*_*z*_), can be extracted by fitting Eq. 1 to the simulation data in Fig. 2c (see Fig. S6 for the fitting procedure). The numerator, *α* (*σ*_*ψ*_, *V*_*z*_), increases with swimming speed but decreases with turbulence intensity, thus encapsulating the aforementioned tug-of-war between turbulence and feeding behavior. The exponent *β* set the degree of nonlinearity of the density-dependent encounter rates in Eq. 1. Overall, *β* could be approximated to *β* ≃ 1.0 (Fig. S6b), with deviations towards smaller values occurring only when the community is well-mixed (high *σ*_*ψ*_ and low *V*_*z*_), a regime in which *α* is negligible and, thus, *E →* 0.

Because, instead of making heuristic arguments, we extracted the encounter rate from our simulation data, Eq. 1 brings a substantial improvement over previous theoretical proposals [28]. Moreover, Eq. 1 allowed us to identify how zooplankton encounters with prey are affected by the environment (i.e. turbulence and prey density) and feeding behavior (i.e. zooplankton swimming speed), key pieces of mechanistic information that allow us to derive next the population-level consequences of microscale patchiness.

### B. Coarse-graining: From micro to macro

The high computational cost of the IBM makes it suitable only for small systems (on the order of decimeters) and, therefore, it cannot be used to address the kilometer-scale plankton patterns observed in the ocean. Thus, to study the macroscale consequences of the microscale patterns above and of their underlying population-level interactions, we implemented a coarse-graining method.

The method exploits the idea from Renormalization Theory [41, 42] that only key components of the microscale dynamics remain relevant at coarser scales and thus are responsible for the resulting large-scale phenomena, as the effects of irrelevant components “average out” at such scales. In other words, a coarser description obtained by looking at the microscale dynamics at larger scales (by, for example, calculating densities at a larger spatial unit of observation) will ultimately only retain the key components of those dynamics. Thus, to obtain the density-field description, we first considered a large square system of linear size ℒ composed of smaller sections (pixels) of size *L* that were computationally suitable for the IBM. Each pixel was assigned a coordinate (*m, n*) within the larger lattice. *P* (*m, n*) and *Z*(*m, n*) denote respectively the number of prey and zooplankton individuals within pixel (*m, n*), while *p*_*m,n*_ ≡ *P*_*m,n*_ (*t*)*/L*^2^ and *z*_*m,n*_ ≡ *Z*_*m,n*_ (*t*)*/L*^2^ represent their corresponding densities. Thus, instead of tracking individuals within each (IBM-friendly) pixel, we calculated the corresponding pixel-specific densities.

Under this coarse description, we followed a computational approach that borrows concepts from probabilistic simulation methods [43–45] to understand the dynamics of such densities. Specifically, we tracked the changes in species density within each time interval, Δ*t*, for different turbulence-velocity scenarios, (*σ*_*ψ*_, *V*_*z*_). In addition to the IBM dynamics within each pixel, individuals can also move between adjacent pixels (either due to flow or behavior, following the IBM rules). Thus, we measured *p* and *z* at the focal and adjacent (top, bottom, left and right) pixels at the beginning of the time interval, with Δ*p*_+_ (*t*) and Δ*p*_−_ (*t*)representing increases and decreases in prey abundance within the time interval and Δ*z*_+_(*t*) and Δ*z*_−_(*t*) those for zooplankton. See *Materials and methods* for further details.

Because we tracked the fate of each individual in the whole system, we could keep track of the changes due to birth-death processes (B) and those due to transport (T), i.e. 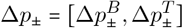 and 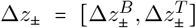. We then defined the rate at which density changes due to the former as 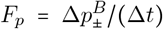 and 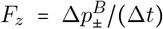, and the latter as 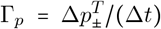 and 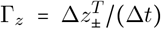. Importantly, the timescale separation between spatial movement and demographic events further resulted in a Markovian property of the coarse representation such that the changes in the densities of a given pixel, (Δ*p*_±_, Δ*z*_±_), could be determined by its current densities and those of its adjacent pixels [46]. For pixels large enough for microscale patchiness to emerge (i.e. *L* > 10 cm; see Fig. S7), these changes in density obeyed Poisson statistics (Fig. S8a,b) with well-defined size scaling laws (Fig. S8c,d) and mean determined by species densities, traits, and environmental conditions (Fig. S8e-h). Altogether, we obtained a density field description for prey and zooplankton at time *t* and at every location, 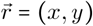, within the larger system, where *x = m · L* and *y = n · L* translate from pixel integer coordinates to real location within the ℒ system:

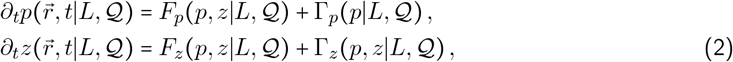

These dynamics depend on the IBM parameters 𝒬 ≡ {*σ*_*ψ*_, *V*_*Z*_, …} and pixel size *L*, the latter setting the spatial resolution of the description (see *Materials and methods*).

Macroscale spatial patterns are typically monitored at pixel resolutions for which the IBM is computationally intractable (e.g. for phytoplankton patterns in the oceans, resolutions range from 120m for the high-resolution satellite images from the SeaHawk Cube Sat mission [6] to 1km for the well-known MODIS database [7]. Therefore, in order to understand what our theory would predict when utilizing (or comparing with) available remote-sensing data, we needed to derive Eq. 2 explicitly for large pixel sizes. To this end, we simulated the IBM for several sizes up to the upper computational limit (*L =* 100cm) and used these simulation data to extrapolate the behavior to larger pixel sizes (see *Supplementary Materials*). This extrapolation was enabled by the well-behaved statistics described above (Fig. S8a,b). Ultimately, for pixels of sizes comparable to the typical resolution of empirical data, this procedure yielded the following expressions:

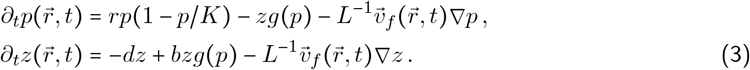

The first term of each equation represents the reproduction and death of prey and zooplankton, respectively; these logistic and linear forms, derived using data from our mechanistic IBM, match the terms typically assumed phenomenologically in population-level feeding models. The last term in each equation accounts for the advection (i.e. directional movement) of individuals due to the flow at the scale of the pixel, which moves around the planktonic predator and prey at the same velocity as the turbulent flow associated with eddies, 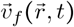 (see *Materials and methods* Eq. 8). An additional feeding-related directional movement also exists but becomes negligible at the kilometer scale when compared to the advection caused by the flow; its demographic effects, however, remain present for all *L*, leading to the second term in both equations, where the density-dependent function *g* encapsulates the microscale patterns induced by active feeding.

As explained in *Supplementary Materials*, further steps in our coarse-graining procedure showed that the feeding function depends on the encounter index *E* and the capture radius *R*_*c*_; specifically, 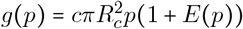. This density-dependent term, which is different from the Holling type I or II functional responses typically used to represent feeding in phenomenological models [47], has substantial consequences for the stability of the community dynamics. For instance, when *E* = 0 (i.e. well-mixed scenario and/or passive zooplankton), the ingestion rate vanishes as *p* → 0, which allows the recovery of the local prey population. In contrast, when *E* > 0 (i.e. active zooplankton) feeding behavior compensates for low prey availability, which leads to high ingestion rates, prey population declines and, potentially, extinction.

To exemplify that such different behaviors can have substantial dynamical consequences, we can look at the mean-field limit (*L* → ∞) of Eq. 3. When feeding is passive, we recapitulate the results of the Lotka-Volterra model (Fig. S9a). However, when feeding is active, trajectories in the (*p, z*) space do not match the typical expectations for this classic model (compare panels b and c to panel a in Fig. S9). Specifically, for active feeding, we found neither periodic oscillatory dynamics when the carrying capacity was assumed infinite, nor inward spirals (i.e. damped oscillations that lead to stationary values) when the carrying capacity was assumed finite. Instead, the data-derived density-dependent interaction term created outward spirals (i.e. unstable oscillations) that can ultimately lead to the extinction of the community (see also linear stability analysis in Fig. S9d-e).

In summary, passive feeding leads to qualitatively different population-level dynamics than active feeding, which might ultimately impact the spatial organization of the community at the macroscale. All these results set us up to tackle our original questions: whether and how macroscopic spatial data can be used to identify behavior across space and time in realistic scenarios. With this in mind, in the next section we adapt Eq. 3 to study macroscale prey pattern formation.

### C. Macroscale dynamics

To study whether zooplankton behavioral signatures can be identified in a dynamic realistic scenario, we simulated upwelling events in the oceans, during which surges of nutrients lead to a wide range of densities for prey targeted by zooplankton. To this end, we modified Eqs. 3 in two ways. First, we added a point-vertex model (see *Materials and methods*) that set the net flow velocity for each pixel, 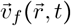, to account for the effects of large-scale eddies in the ocean (see Fig. S1); the angular speed of the eddies was controlled by the intensity parameter, *ψ*. Second, we introduced spatially explicit dynamics for the prey carrying capacity, 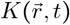, to model the heterogeneity of a growth-limiting nutrient that sets the maximum population density in the absence of predation. Specifically, the carrying capacity followed the equation:

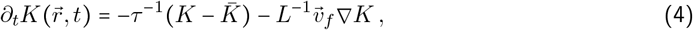

where 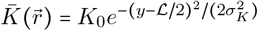 mimics a continuous input of nutrients occurring during the upwelling event that, if not depleted or perturbed by consumers, forms a horizontal band of width *σ*_*K*_ in the center of a ℒ = 100-km squared domain (Fig. S10a). The parameter *τ* represents the typical time for any initial condition for 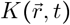 to converge to the horizontal band 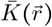 in the absence of turbulence; we set *τ* = 8 days, per [31]. For negligible eddy effects, the velocity of the flow is *v*_*f*_ = 0 and thus 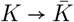, but *K* will deviate from the imposed profile for moderate and strong eddies (Fig. S10b and first panel of Fig. 3a). We initialized this dynamic carrying capacity as 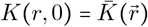 with the aim of understanding how prey dynamics and turbulence disrupt such a marked initial pattern (the band); for robustness, we also explored other initial conditions and carrying capacity profiles (Fig. S10c,d), which did not alter our conclusions (see below).

**Figure 3:**
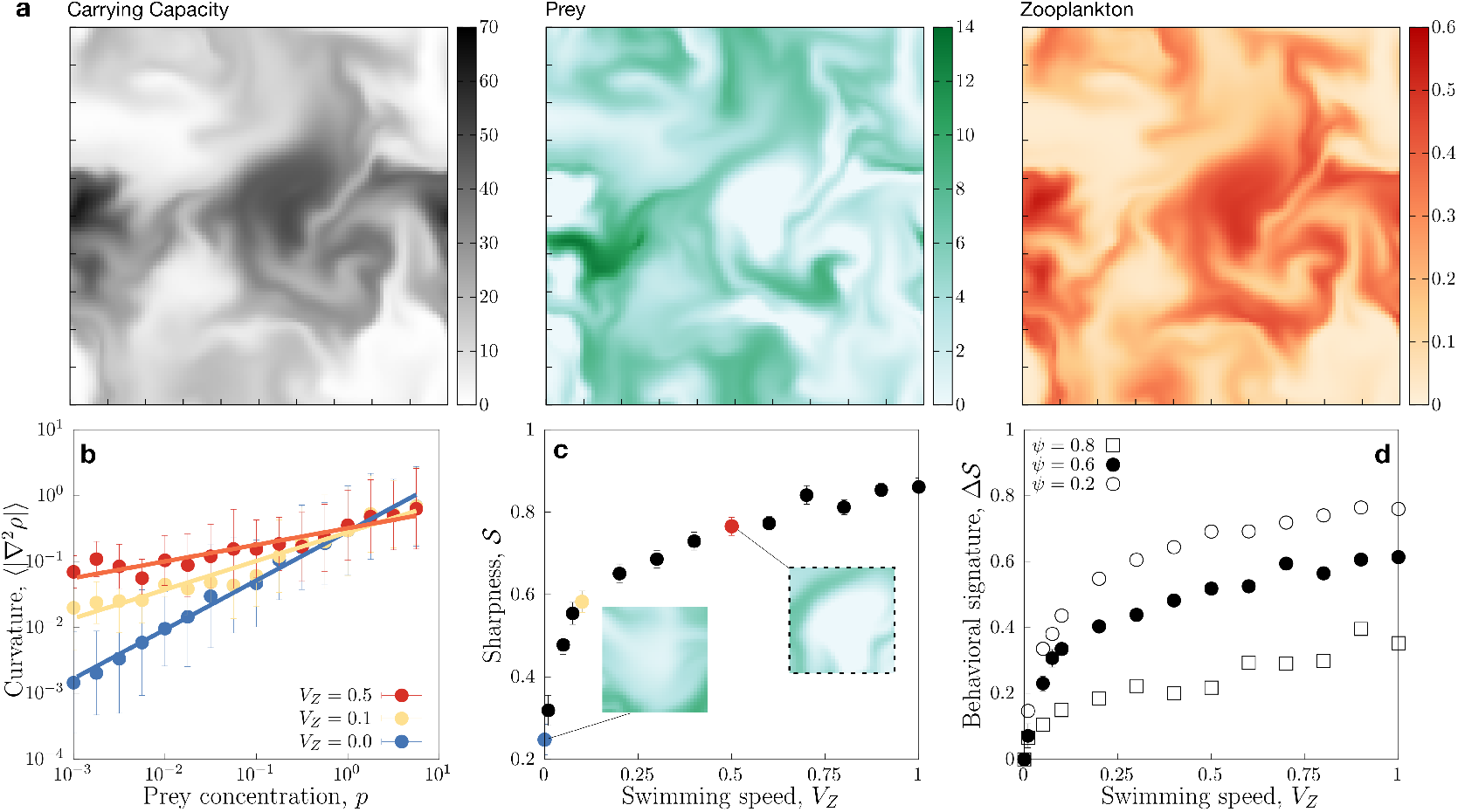
Zooplankton behavioral signatures in macroscale patterns of prey. a) Spatial patterns for carrying capacity, prey, and zooplankton (respectively, from left to right) in our simulated upwelling event. Color indicates densities in ind./ml. b) Curvature-density relationship for different swimming speeds, *V*_*z*_. c) Pattern sharpness as a function of *V*_*z*_. Insets highlight typical boundaries for *V*_*z*_ = 0.0 cm/s and *V*_*z*_ = 0.5 cm/s (the latter corresponding to the dashed square in the prey pattern from panel (a)). d) Behavioral signature, defined as difference in sharpness relative to the passive case, Δ𝒮 ≡ 𝒮(*V*_*z*_) − 𝒮(0), for different turbulence levels, *ψ*. For these simulations, the total domain size was ℒ = 12 km (100 cells of size *L* = 120 m) and a horizontal carrying capacity band, 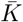, of length 12Km, width *σ*_*K*_ = 840 m, and amplitude *K*_0_ = 100 was placed around the middle of the vertical axis. We adjusted eddies to introduce a mean flow speed 1km/day (*ψ* = 0.6 day ^1^) and fixed *σ*_*ψ*_ = 0.25 cm/s to keep the diffusion and advective timescale comparable to experimental values [31, 39]. The snapshots used for these figures were taken close to the peak of prey density during the upwelling event (approx. *t =* 12 days, as seen in Fig. S11a). We initialized the system using low, homogeneous prey and zooplankton densities (1 and 10^−2^ ind./cm^2^, respectively).

With this setup, we integrated numerically our complete large-scale model following standard schemes (see *Supplementary Materials*) and studied how the spatio-temporal prey pattern depends on predator swimming speed, with the goal of discerning signatures of feeding behavior. In all cases, we initialized both zooplankton and prey populations using small densities. Regardless of the (fixed) *V*_*z*_ value, the initially negligible top-down regulation allowed prey to grow rapidly to a peak of high density (Fig. S11a and prey panel in Fig. 3a). The prey surge then facilitated the growth of predator density (zooplankton panel in Fig. 3a), which consequently led to a decrease in prey density (Fig. S11a). Consistent with our mean-field findings, passive feeding led to stable coexistence of prey and predators, whereas active feeding ultimately led to the extinction of prey.

As this realistic and expected sequence of events unfolded, zooplankton dynamically shaped the emerging prey patterns. Importantly, the wide range of prey densities produced in space and time by these events offered an opportunity to investigate whether the density-dependent feeding effects described in the previous section were present in these macroscale spatial patterns. Based on our results in Fig. 2c, we hypothesized that behavioral signatures emerge and should become more evident closer to the expanding edges, where prey density is lower. Specifically, we predicted that active feeding should produce a more noticeable contrast between the interior of the prey patterns and the borders (i.e. sharper boundaries) than passive feeding. Although visually that seemed to be the case (see e.g., Fig. S11b), we also quantified the morphological differences caused by different swimming speeds by measuring the dependence between the curvature of the pattern and prey density [48]:

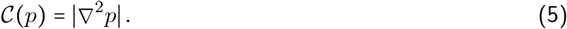

The function 𝒞(*p*) provides a proxy for the curvature of the prey density field *p* because, for a given location 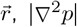 quantifies density differences in any direction of the two-dimensional system. For smooth patterns, curvature vanishes as prey density asymptotically goes to zero. For sharp boundaries, in contrast, prey density abruptly drops to zero and, consequently, the curvature takes a non-zero value (see *Materials and methods* for further details and examples).

Using Eq. 5, we measured curvature at any given time and every location in our simulated patterns, and extracted a relationship between curvature and prey density. We observed that this relationship approximated a power-law throughout the duration of the upwelling event, although its slope varied with time (compare *t* = 2, 12, 14 in Fig. S11c). This power law emerged from the interaction with zooplankton and, at any given time, its slope reflected their feeding behavior (i.e. the magnitude of *V*_*z*_; see Fig. 3b and Fig. S11c). In particular, the slope captured clearly the difference between active and passive predators, which we traced to differences in swimming behavior rather than in overall increase in ingestion (passive zooplankton with a higher ingestion rate cannot induce similar signatures, see Fig. S11d). In other words, this finding confirmed our hypothesis that feeding induces behavioral signatures in prey macroscale patterns, and that these signatures can be inferred by measuring the curvature of boundaries or, more specifically, the difference in the slopes. Thus, to quantify the sharpness 𝒮 of the boundary at a given time we used the slope of the curvature-density relationship: log⟨𝒞 (*p*)⟨ ~ (1 − 𝒮) log *p* where (𝒞(*p*) is the pattern curvature averaged over pixels that show a given prey density, *p*.

With the expression above, we could infer a sharpness for any given swimming speed, 𝒮 = 𝒮 *V*_*z*_ (see slopes in Fig. 3b and summary panel for S as a function of *V*_z_ and time in Fig. S11e). An ideal observation period encompasses the peak of prey density during the upwelling event because it exhibits the widest range of densities that result from model dynamics. We indeed observed fingerprints of feeding behavior in sharpness (Fig. 3c), as 𝒮 showed a very low value for simulations with passive zooplankton (*V*_*z*_ = 0), and increased smoothly but steeply when we instead considered active zooplankton (*V*_*z*_ > 0). A similar phenomenology was observed at other times across the duration of the upwelling event (Fig. S11e). Therefore, sharpness reflects behavioral signatures, and this abrupt change in value provides a way to discern between passive and active predators. The ability to notice these differences depended on turbulence: as one might expect, at high turbulence (large *ψ*) the strong eddies blurred behavioral signatures (Fig. 3d).

#### Behavioral diversity can induce spatial niche partitioning

To test our behavioral-segregation hypothesis, we extended our model—which, for simplicity, considered that all zooplankton had the same, fixed feeding behavior, *V*_*z*_—to allow for behavioral diversity. Zooplankton communities typically show diversity in feeding behavior [28], with some species even showing adaptive (i.e. context-dependent) feeding [49, 50]. There are alternative approaches to making the simplest modeling extension that allows for behavioral diversity, and here we explored two of them for robustness: (i) we assumed two co-occurring species of zooplankton, one passive and one active (see *Supplementary Materials* and Fig. S13); alternatively (ii) we assumed a single species of zooplankton able to adapt its feeding behavior, i.e. change from passive (*V*_*z*_ = 0) to active (*V*_*z*_ > 0) feeding depending on metabolic costs and benefits [51].

Below, we focus on (ii) and modify Eq. 3 such that the feeding function changes dynamically reflecting the differences between the metabolically costly but informed swimming associated with active feeding, and metabolically cheap “sit-and-wait” feeding associated with passive feeding that relies on random encounters (see Eqs. 9 and 10 in *Materials and methods*). In this extended version of the model, at each point in space and time zooplankton show the feeding behavior that yields the higher ratio between ingestion and metabolic cost (i.e. higher fitness; see *Materials and methods*). As a result, passive feeding is beneficial at high prey densities while active feeding is beneficial at lower prey densities (Fig.4a). Because velocity is an essential component differentiating feeding behavior (and thus the feeding function), the emergent prey density threshold that determines whether active or passive feeding is advantageous is velocity-dependent (Fig. 4a). Fig. 4b shows a snapshot of the prey density obtained with the adaptive-feeding model at the peak of an upwelling event, while Fig. 4c shows the behavior of the predator population in the same snapshot. Together, Fig. 4a-c panels confirm that, when accounting for the main energetic costs of feeding even in this simplistic way, zooplankton switch behavior dynamically, with active behavior dominating at low prey densities and passive behavior dominating at high densities.

**Figure 4:**
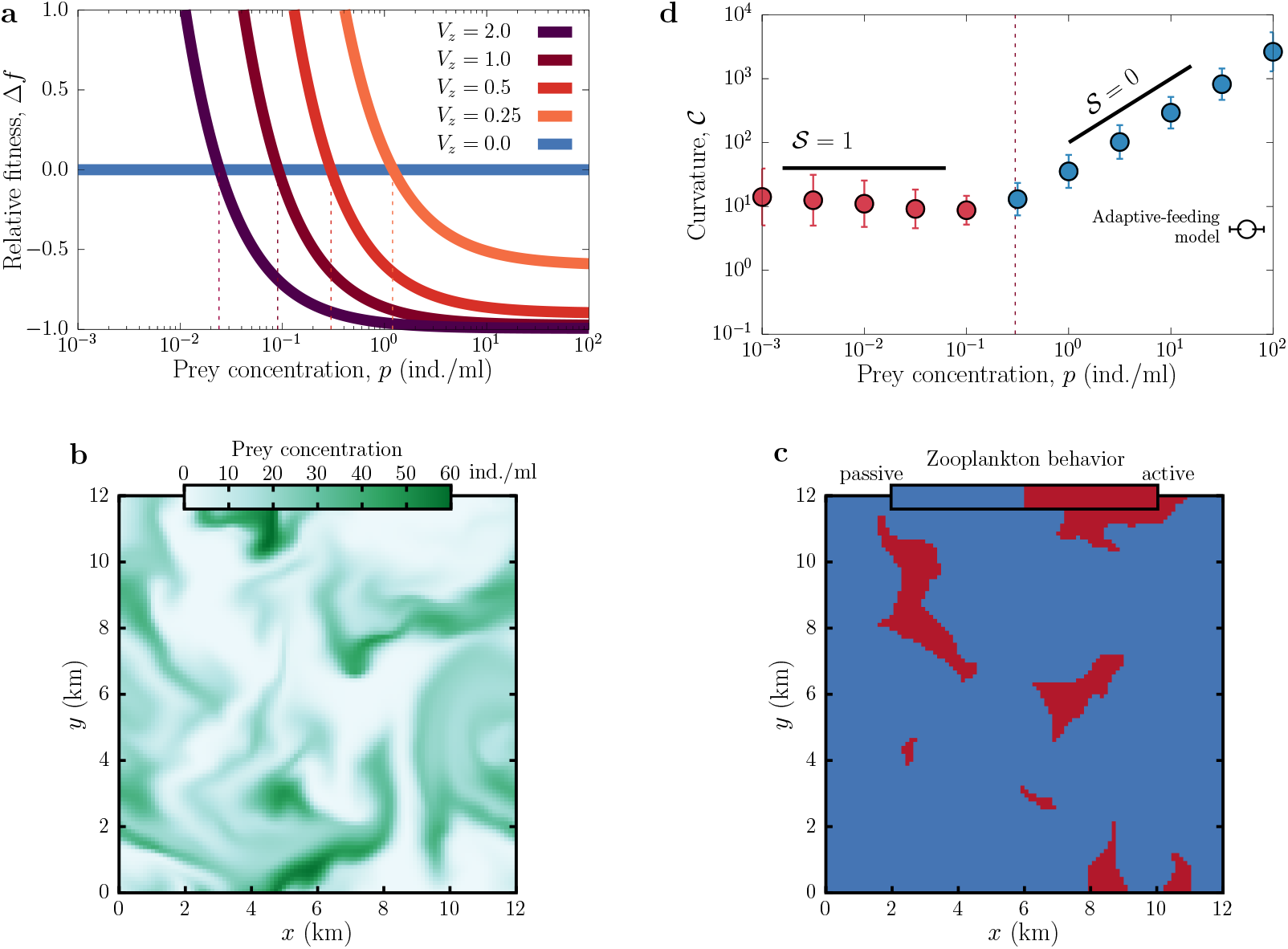
Spatial niche partitioning. Anti-clockwise: a) Relative _tness di_erence, Δ*f* = (*f*_1_ − *f*_0_)/*f*_0_, for different swimming speeds taking the passive case (*V*_*z*_ = 0:0 cm/s) as reference (i.e. *f*_0_ and *f*_1_ corresponds to fitness for passive and active cases respectively). Vertical dashed lines highlight, for each swimming speed, the prey density threshold below which being active is advantageous (Δ*f* > 0). b) Prey density and c) zooplankton behavior patterns for the adaptive-feeding model with *V*_*z*_ = 0:5 cm/s, for a snapshot close to the peak of the upwelling event. To match realistic concentrations, we set *K*_0_ = 10^5^ and remaining parameters as in Fig. 3a. d) Curvature-density relationship extracted from our adaptive-feeding model with *V*_*z*_ = 0:5 cm/s. The curve is an average over 50 snapshots (to account for the different sources of variability such as initial condition and eddy distribution) taken within a 4-day time-window right after the upwelling event; the vertical line indicates the density threshold identi_ed in panel a for *V*_*z*_ = 0:5, and dot color indicates the dominant behavior for each prey density (see colorbar in panel c). For all panels, *q* = 4, *d*_1_ = *d*_*0*_ = 0:10 day^−1^, *c*_0_ = 10.

Such behavioral segregation can indeed be inferred from the shape of prey patterns just by monitoring sharpness, 𝒮 (*p*), which transitioned from low (𝒮 ≈ 0) to high (𝒮 ≈ 1) values as prey density decreased (Fig. 4d). Sensitivity analyses using different initial conditions for the nutrient input (band versus hot-spots) confirmed that this shift in sharpness is robust for high and low concentration regions (Fig. S12), which reinforces the robustness of our main (qualitative) takeaway. Interestingly, the threshold density where the transition from passive to active occurs is quite sensitive to initial conditions as well as to the parameters representing the costs and benefits of active vs passive feeding.

The robustness of our findings was further reinforced by the fact that we obtained similar qualitative results under a scenario of behavioral diversity where we extended the original model to include an only-passive zooplankton species and an only-active species (scenario (i) above, see Fig. S13). The two species were able to coexist precisely because of an emergent spatial niche partitioning, whereby the passive species outcompeted the active one at high prey densities, but was outcompeted by the active one at low prey densities. However, it is worth noting that these competitive outcomes are most definitive at extreme values of prey density; in a broad intermediate region, the theoretical outcome is blurred by elements of the dynamics, such as turbulence replenishing prey locally in an area where active zooplankton were just beginning to grow and assert their dominance. Although studying such dynamics is beyond the scope of this section (where we simply want to establish the qualitative existence of the two behavioral regimes), they are interesting avenues for future work, especially work that tries to study the specific details of a diverse zooplankton community. Nonetheless, and although the reality of oceanic patterns is much more complex than our models, these results overall support the hypothesis that zooplankton behavior might contribute to shaping prey spatial patterns.

### D. Conclusions

Our multiscale framework provides a previously computationally inaccessible bridge across scales, from the micro- and millimeter-scale of individuals to the centimeter- and meter-scale of species interactions, and ultimately to the kilometer-scale of macropatterns. Built on a mechanistic individual-based model and scaled via a rigorous coarse-graining procedure, our framework is tractable without having to sacrifice crucial biological detail. This allowed us to understand how prey spatial patterns at different scales may be regulated by zooplankton behavioral traits.

At the microscale, we found that patchiness is characterized by two emergent spatial features: an increase in interspecific encounters and an aggregation of zooplankton, both of which are regulated by turbulence, predator swimming speed, and prey density. These findings bring mechanistic insight to existing empirical work. Our work provides theoretical support for the field and experimental studies on the impact of turbulence on zooplankton feeding behavior and capture success, which have shown that encounters with prey are affected positively by active swimming and negatively by turbulence [28, 52, 53]. Additionally, previous work observed that reproducing the different capture success at low densities of passive and active feeding behavior requires the use of qualitatively different feeding response functions [49, 54]; in our framework, such functions emerged from the analysis of our IBM, thus allowing us to trace mechanistically their origin. Further, our framework shows that the observed small-scale zooplankton aggregation and its negative dependence on turbulence (observed via towed video microscopy [55]) can be explained simply from emergent, indirect correlations between zooplankton individuals in active search and feeding of prey; previous models assumed ballistic or random zooplankton swimming behavior, and thus could not reproduce this pattern, or they did so only after invoking attractive social forces between zooplankton individuals. (e.g. [27]).

Our coarse-graining method allowed us to translate this microscale emergent pattern into effective population-level interaction rates; by stress-testing the IBM under different scenarios, we obtained a density field description that was fully parameterized by species traits and the physical environment. Analyzing simulation data statistically to infer population-level expressions thus enabled a rigorous mapping that connected scales. Previous attempts to connect individual and population levels either estimated population-level encounter rates by, for example, resorting to heuristic, geometry-based, arguments [28]; calculated feeding flow using hydrodynamical models [33, 56]; or developed analytical approximations to account for attraction-repulsion between individuals [27]. However, in these approaches the effects of collective emergent phenomena (e.g., density-dependent encounters) were not accounted for, mostly due to the intrinsic complications when dealing with spatially-extended systems [57, 58]. Thus, existing models could not provide a mechanistic link between individual and population levels. Here, we used computational methods to overcome analytical limitations and aid us in the construction of dynamic equations based on our simulated data [59, 60]. Because, according to Renormalization Theory, “irrelevant” details are lost during coarsening, these equations represent the fundamental mechanisms underlying the interactions between zooplankton and their prey, with density-dependent feeding playing a particularly relevant role in the emergent large-scale pattern.

Overcoming this theoretical “problem of scale” allowed us to access valuable ecological information about plankton communities across scales. In particular, we were able to map the curvature of macroscale prey patterns to the swimming speed of the zooplankton that is harvesting it, which revealed spatial changes in dominant zooplankton behavior driven by the relative costs and benefits of passive versus active feeding. Our approach thus provides what, to our knowledge, is the first bottom-up framework to identify predator behavior from macroscale prey patterns. Establishing a two-way connection across scales could, for example, be used to monitor which zooplankton (or, at least, which feeding behavior) dominate during the different stages of an upwelling event at given spatial locations. This information can be helpful for understanding not only the dynamics of the microbial loop and oceanic biogeochemistry, but also any associated ecosystem services (e.g., fisheries with preference for particular zooplankton types). Note that, although nutrients and other sources of regulation, such as viruses, are also expected to affect the shape of oceanic macroscopic patterns such as those associated with upwelling events, the robustness of our results against e.g., different nutrient spatial profiles emphasizes the importance of feeding behavior as a key driving factor for the curvature of the pattern.

We focused here on the perception and active swimming of a generic zooplankton to demonstrate the power of a framework that accounts for individual-level detail; in the future, extensions of our framework could easily be made to incorporate more specific aspects of feeding that have been shown to be important for pattern formation at small scales, such as social interactions [27] and gyrotaxis [23]. A further developed version of our framework that captures how behavior varies across taxa could explain mechanistically patterns of structural variability that have been recently cataloged [10]. Because the expectation is that changes in environmental conditions will alter feeding behavior [49], our framework provides a diagnostic (if not predictive) tool to identify those behavioral changes through the analysis of macroscale patterns. For example, climatic changes affecting zooplankton physiologically and/or the strength of turbulence will alter the costs and benefits of active feeding: if, for instance, short- or long-term climatic changes increase turbulence, higher mixing will make the investment in active feeding less beneficial for zooplankton, and therefore curvature will transition from non-zero to zero slope at smaller *p* values. This means that sharp edges will be found only in more prey-depleted areas of a pattern, and therefore overall patterns will look more generically smooth. Nonetheless, although our methodology suggests a non-invasive way to make inferences regarding prey and zooplankton interactions and behavior (that is, by measuring the sharpness of remote-sensing images of prey macroscopic patterns), the validation of such inferences will still require a combination of such remote-sensing images with in-situ data (for example, to discern the extent to which passive feeding results from zooplankton versus, e.g. viruses).

Beyond the marine context, our computational multiscale framework can be easily applied to and be useful when analytical and computational approaches to scale up individual-level behavior are challenging: for example, when organisms are moving based on complex decision-making strategies [61] that account for the different costs and benefits of individual actions (e.g., fish schools and starling flocks aiming to avoid predatory attacks) [62]. For such cases, coupling with our coarsening method either a mechanistic IBM tailored to the focal system or empirical individual-level data enables an exact and mechanistic connection between scales. This approach provides an alternative to the strong mathematical barriers that arise in analytical methods [27, 57], with which to check for potential behavioral signatures in large-scale patterns.

## II. MATERIALS AND METHODS

### Individual-based model description and parameterization

Individuals exist within a square domain of size *L* with periodic boundary conditions (i.e. torus topology). The individual-based model (IBM) accounts for prey and zooplankton birth-death processes, zooplankton feeding behavior, and the influence of an external flow. We implemented these processes using a time-dependent Gillespie algorithm [63] (see *Supplementary Materials* for detailed description) We parameterized our model using values that are generically applicable to zooplankton and their prey. For the former, we aimed to capture the range of behaviors observed from microzooplankton to fish larvae, which is therefore also reflected in the choice of parameter values for the predator and the prey (see Table 1). Thus, we set the prey reproduction rate *r* = 1 day^−1^ and the zooplankton death rate *d* = 0.1 day^−1^. For computational convenience, we set the prey carrying capacity at *K* = 100 individuals/ml. Zooplankton catching range, *R*_*c*_, and perceptual range, *R*_*p*_, were set as a function of body size (i.e. the effective radius, set to *B*_g_ = 0.2 cm for our generic zooplankton [28]) as *R*_*c*_ = *B*_*g*_, *R*_*p*_ = 5*B*_*g*_. For the chosen body size, zooplankton velocities can reach values around *V*_*z*_ ~ 0.5 cm/s [28], which will be our ‘focal’ feeding speed for active zooplankton. With this parameterization and considering a search rate *c* = 30, depending on prey availability, ingestion rates ranged from a couple of prey per minute to a prey individual every few seconds, which is within observations (see experimental reference values in, e.g., [64, 65]). For more details see Fig. S2.

### Spatial characterization of the microscale patchiness

To quantify the interspecific correlations of the emergent patterns, we monitored through time the number of prey that were accessible to each zooplankton individual (i.e. within its catching range, *R*_*c*_). For a fixed pair of prey and predator densities, (*p, z*), the per-predator average number of potential targets reached a constant value ⟨*P* | *R*_*c*_⟩ after a short transient (Fig. S4a). Analogously, to quantify predator aggregation we measured the stationary value of the number of zooplankton individuals that were within a focal predator’s catching range, ⟨*Z* | *R*_*c*_ ⟩ (Fig. S4b). Using these two measures, we calculated the encounter index *E*, which quantifies how much prey availability there is per predator, and the clumping index *C*, which quantifies predator aggregation, both relative to the values expected for the well-mixed scenario [27]:

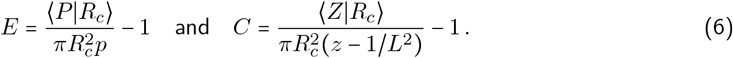

where the denominators represent the expected prey targets and zooplankton individuals, respectively, for the well-mixed case. With the definitions above, if the community is well-mixed (i.e. individuals are randomly distributed in space), *E* = *C* = 0; a positive spatial correlation between prey and zooplankton is indicated by *E* > 0; and correlation among zooplankton, by *C* > 0.

### Point-vortex model

In order to account for the key elements of turbulence at large-scales, we introduced a fixed number of vortices (or eddies) *N*_*e*_ =1000 in our spatial domain. Setting the variability of the size and rotation direction of the eddies in a realistic way leads to a stirring process with scale-free velocity spectrum [31, 66]. We used the following stream function to capture these features:

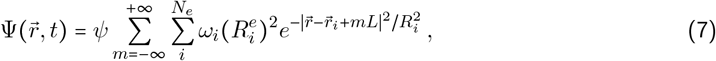

where *ψ* controls the angular speed of eddies (in units of day^−1^), *ω*_*i*_ ∈ {−1, 1} sets the rotation direction, while 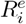 and 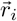 are the radius and center of each eddy *i* (for *i* = 1, 2, …, *N*). Due to the imposed periodic boundary conditions, the influence of the vortices looped around the domain, which was accounted for by the sum over *m*; since this effect decays exponentially fast, however, considering only *m* = −2, −1, …, 2 already achieved a precision 𝒪 (10 ^−30^). The flow velocity, *v*_*f*_, is the curl of the stream function:

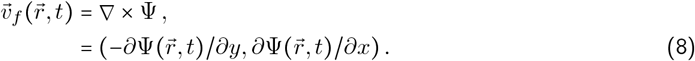

We sampled vortex sizes from a power-law probability distribution, *P* (*R*^*e*^) = (*R*^*e*^) ^−3^ (for *R*^*e*^ ∈ [ℒ/40, ℒ/4]) and obtained a scale-free transversal velocity spectrum ∝ *k*^−3^, where *k* is the spatial frequency (Fig. S1). This probability distribution generated a power-law spectrum with exponent −3, characteristic of geostrophic turbulence occurring at large scales [31].

### Adaptive-feeding model

The adaptive-feeding model modifies Eq. 3 to account for a metabolically mediated switch between passive and active behavior. Specifically, the model introduces two modifications that aim to reflect the costs and benefits of each of the two feeding strategies.

The first change consists in replacing the feeding function, *g* (*p*), by:

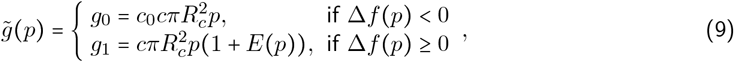

where *c*_0_ > 1 reflects the reduced energy expenditure and reduced risk of feeding-related predation associated with passive feeding [51, 67]. To assess the sensitivity of our findings to this new parameter, we explored a broad range for *c*_0_ ∈ [2, 100], which can result from a variety of factors; for example, passive zooplankton are 2-8 times less likely to be predated upon than active zooplankton [68], which motivated the moderate value *c*_0_ = 10 chosen for illustration purposes in Fig. 4d [69].

The second change consists in modifying the mortality rate *d* (which, since the term reduces growth, can be seen also as a proxy for metabolic costs) for active predators:

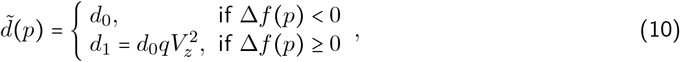

where *q* is a fixed hydromechanical factor and 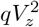 reflects the energy dissipated due to drag, consistent with Stokes’ Law [51].

The behavioral switch is controlled by the relative difference in associated “fitness” Δ*f* (*p*) = (*f*_1_ − *f*_0_)/*f*_0_, where *f*_0_ = *g*_0_/*d*_0_ and *f*_1_ = *g*_1_/*d*_1_ are simply defined as the ratio between the benefits and costs of the corresponding behavior [51].

Altogether, these modifications extend our previous model by introducing a trade-off between the two behaviors: the energy-saving but potentially unproductive passive behavior versus the expensive but targeted active behavior.

### Measuring the curvature-density relationship

We selected hourly snapshots following the peak of prey density during the upwelling event. Immediately after the peak is when the widest range of prey densities are observed, and feeding starts significantly shaping the (very abundant) prey community. For these reasons, and although the curvature curves looked approximately as power laws at any time along the duration of the upwelling event, our analysis of the sharpness of the prey pattern focused on averages obtained within 4 days following the peak.

For each snapshot of the system, we obtained the associated local curvature and density for every pixel; we used a log (base 10) scale, [log|∇ ^2^*p* |, log *p*], to better elucidate a potential linear (or non-linear) relationship. We then binned the data according to prey density, and obtained the expected value of the logarithm of the curvature within each bin, generating the curve ⟨log|∇ ^2^*p*|⟩ vs. *p* (Fig. S11c).

For smooth patterns, the profile becomes flat as prey density goes to zero at the borders, and thus the curvature 𝒞 vanishes; for example, if we had an ideal circular pattern and the boundary of the pattern were described by the smooth function *p* = *e* ^*−r*^, where *r* represents the distance to the center of the pattern, the associated curvature would be 𝒞(*p*) = |∇^2^*p*| ~ *p*, which vanishes as *p* → 0, exhibiting zero sharpness, 𝒮 = 0. For sharp boundaries, in contrast, the density profile drops abruptly to zero, and concurrently so does the curvature to a non-zero value; as an example, a sharp radial density profile given by *p*(*r*) = 1 − *r*^2^ if *r* < 1 and *p* = 0 if *r* > 1 leads to a constant curvature 𝒞 = |∇^2^*p*| = 2 > 0 yielding 𝒮 =1 even at the pattern boundary, where *p* → 0.

## III. CODE AVAILABILITY

The code to reproduce all results and run all simulations, as well as the python script for the extraction of the curvature-density relation are available at https://github.com/ehcolombo/SignaturesPP.

## IV. ACKNOWLEDGMENTS

We thank Ron Shvartsman and Laure Resplandy for their critical, in-depth reading and constructive feedback on the manuscript. We are grateful for feedback from members of the Tarnita and Bonachela labs. We acknowledge support from the Gordon and Betty Moore Foundation (#7800; E.H.C, C.E.T., J.A.B.), the Simons Foundation (#82610; J.A.B) and NSF DMS-2052616 (J.A.B.).

## Notes

### Competing Interest Statement

The authors have declared no competing interest.

