## Supplementary materials for "Zooplankton feeding behavioral signatures in the morphology of macroscale prey spatial distribution"

Eduardo H. Colombo,[1,2,3,*] Corina E. Tarnita,[1,4] and Juan A. Bonachela[2,†]

[1] *Department of Ecology & Evolutionary Biology, Princeton University, Princeton, NJ 08544, USA*
[2] *Department of Ecology, Evolution, and Natural Resources, Rutgers University, New Brunswick, NJ 08901, USA*
[3] *Present address: Center for Advanced Systems Understanding (CASUS), Helmholtz-Zentrum Dresden Rossendorf (HZDR), Görlitz, Germany.*
[4] *Department of Environmental Systems Sciences, ETH Zürich, Zurich, Switzerland*

### A. Individual-based model description and implementation

We implemented the ecological processes in our individual-based model using a time-dependent Gillespie algorithm as follows. Note that we refer to zooplankton indistinctly as predator henceforth.

*i) Demographic events*, namely reproduction, death, or zooplankton feeding of a prey individual, occur stochastically at rates that are state-dependent, that is, depend on the density of prey and zooplankton and, therefore, change with time (see [1] for a similar implementation). To account for this time dependence, time intervals need to be calculated for the algorithm by normalizing by such rates. First, for every time step (duration $\delta t = 0.1$s), we calculate the instantaneous characteristic time to a demographic event, defined as the inverse of the sum of all demographic rates, $\tau = [rP(1 - P/(L^2 \cdot K) + Zd + \sum_{j=0}^{Z} c\gamma_j]^{-1}$, where $r$ is the prey reproduction rate, $K$ is the prey carrying capacity, $d$, the zooplankton mortality rate, and $\gamma_j$ is the number of prey individuals within reach for predator $j$ (see symbols, values, and units in table I). Thus, we can define the normalized waiting time to the next *demographic* event, $\delta t/\tau$, which can also be seen as the probability for the next event to be a demographic one. We then pick a random number $\tau_1$ exponentially distributed with mean one, representing the normalized waiting time to the next event (of *any* type). If $\tau_1 > \delta t/\tau$, there is no demographic event within the time interval, and thus during $\delta t$ individuals diffuse due to small-scale turbulence (see item ii below) and then swim towards prey (if active zooplankton, see item iii). On the other hand, if $\tau_1 < \delta t/\tau$, a demographic event will occur during the $\delta t$ interval. In this case, individuals diffuse during a time interval $\delta t' = \tau_1 \tau$ to the demographic event; such time interval is then discounted from the time to the next target-selection-and-swimming event , $\tau_J - \delta t'$. The type of demographic event that occurs after $\delta t'$ is chosen at random with a probability calculated as their relative rate, e.g., feeding events have a probability $czg(p)L^2\tau$ to occur. If a natural mortality event occurs, random individuals are removed from the system. If a birth event occurs, a new individual is placed at a random location side-by-side to the parent. If a feeding event occurs, a predator is selected at random with a probability given by the relative abundance of prey within catching range (i.e. prey individuals within catching range divided by total number of prey). We consider the catching step to be instantaneous and thus it is not explicitly implemented, although a version of the IBM with explicit catching did not show qualitative of quantitatively different results. After the occurrence of the demographic event, a new $\tau_1$ is sampled and the algorithm repeats to check whether a new demographic even occurs in the remainder of the interval, $\delta t - \delta t'$. When the full $\delta t = 0.1$ is exhausted, the algorithm is restarted for a new $\delta t = 0.1$ interval.

*ii) Small-scale turbulence:* At the spatial scale that is the focus of our IBM, the flow velocity field can be described by means of Gaussian random fluctuations [2, 3]. We thus defined the components of the flow velocities, $v_x$ and $v_y$, which are stochastic variables drawn from a Gaussian distribution, $p(v) = (\sqrt{2\pi\sigma_\psi^2})^{-1} e^{-|v|^2/(2\sigma_\psi^2)}$, where $v = v_x$ or $= v_y$ and the intensity of the fluctuations, given by $\sigma_\psi$ (all in cms$^{-1}$), is the same in both directions. The position for each individual, $r = (x, y)$, is updated following a standard first-order stochastic algorithm, $x(t + \delta t) = x + \sigma_\psi \xi(t)\sqrt{\delta t}$ (same for $y$), where $\xi$ is a zero mean Gaussian white noise.

*
†

*iii) zooplankton swimming behavior* is implemented in the form of swimming triggered by the presence of prey. Every $\tau_J = \delta t = 0.1\,\text{s}$, all predators evaluate their surroundings (i.e. area within their perception range) and, if the zooplankton individual is active, swim towards a randomly chosen targeted prey individual. The probability for a prey individual to be chosen decays linearly with distance to the predator, and becomes zero exactly at $R_p$. Once a target is selected, the zooplankton individual moves to a new location given by $\vec{r}(t + \delta t) = \vec{r}(t) + V_z \delta t\ \hat{\theta}$, where $V_z$ (in $\text{cm}\,\text{s}^{-1}$) is the swimming speed and $\hat{\theta}$ is the unitary direction vector that points towards the target. If the distance to the new location is larger than the actual distance to the target, the predator stops at the target location. In the case that there are no prey within the perceptual range, behavior is not triggered and thus $\hat{\theta} = 0$. Ultimately, the number of prey in reach $\gamma_i$, which controls the feeding rate, will be not only controlled by prey density itself, but also by swimming speed and turbulence, as they control the correlation between prey and zooplankton. In Fig. ST1, we show the fraction of time predators have prey in reach as a function of prey density for different swimming speeds, highlighting how the main differences between behaviors are noticeable as prey density decreases. The rules above were implemented using C/C++ code with parallel schemes (*OpenMP*).

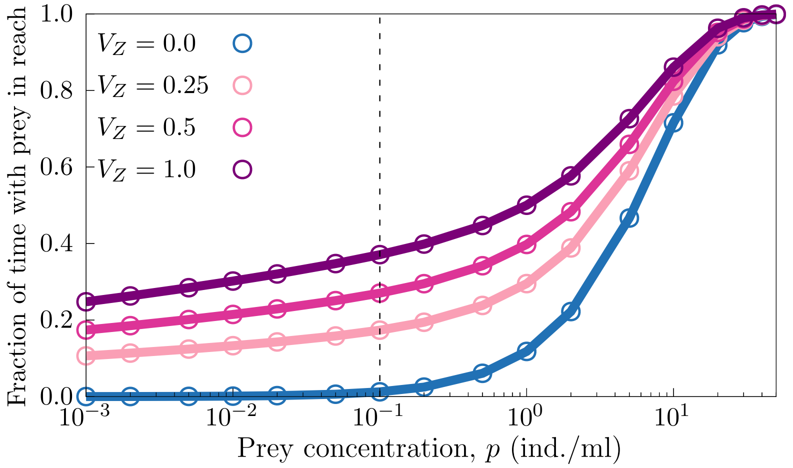


Figure ST1. **Fraction of time with prey in reach.** Solid lines show the fraction of time between ingestion events for which zooplankton have prey in reach but do not capture/ingest them. For the typical speed of an active-like predator, $V_z = 0.5$ cm/s, the fraction of time spent when prey density is low (regime for which feeding behavior matters most and the time between events is longer, see main text) is under 0.1; the fraction is $\simeq 0$ for the passive case. For high prey densities, as expected, prey individuals are within reach at all times regardless of feeding behavior. Fraction of time obtained from data (dots), estimated using the statistics of the number of prey within reach $n_p$ (see Fig. S8), under the assumption that $n_p$ follows a Poisson statistics with mean $E$ given by Eq. 1 in main text. Specifically, the characteristic time-window size within which $n_p > 0$ is $1/(1-q_0)$, and with $n_p = 0$ is $1/q_0$, where $q_0$ is the probability of having $n_p = 0$ prey in reach in a given instant; thus, the fraction of time for which $n_p > 0$ is then $q_0/(1-q_0)$ (solid lines). For these results, $\sigma_\psi = 0.25$ cm/s (remaining parameters as in the main text).

#### B. Toy model to understand optima in encounter index curves

The toy model configuration assumes a one-dimensional domain ($10cm$) with periodic boundary conditions, two prey individuals are kept fixed at locations $-d/2$ and $d/2$ respectively, thus being $d\ cm$ apart from each other. A single active zooplankton individual is considered that swims towards prey with speed $V_z$ while being affected by small-scale turbulence, as in the IBM. This scenario allows us to understand how the encounter index is being maximized at a finite (optimal) speed, $V_z^\star$. Fig. ST2 (left panel) shows the encounter index as a function of swimming speed for different values of $d$. For the case $d < R_c < 2d$ (solid black line), the right panel shows the probability distribution for the predator location for different swimming speeds. Both plots combined show that the maximum encounter index is achieved for $V_z$ for which the predator is expected to stay locked in between the two prey.

For $V_z < V_z^\star$, there was an expected positive correlation between swimming behavior and encounter rates (i.e. swimming faster increased the encounters with prey). For speeds larger than $V_z^\star$, however, zooplankton committed to the signal received just before moving towards the target, and the swift approach to the targeted prey led to missing (or reducing) the chance of opportunistic encounters with non-targeted prey that may have been found on the way. In other words, a tradeoff between speed and resulting encounters emerged, thus leading to the maximum in the encounter index curve. Increased mixing due to higher turbulence reduces the advantage to swimming and thus the disadvantage of missing opportunistic encounters, which is why an increased turbulence leads to the disappearance of the optimal swimming speed (as in in Fig. 2b).

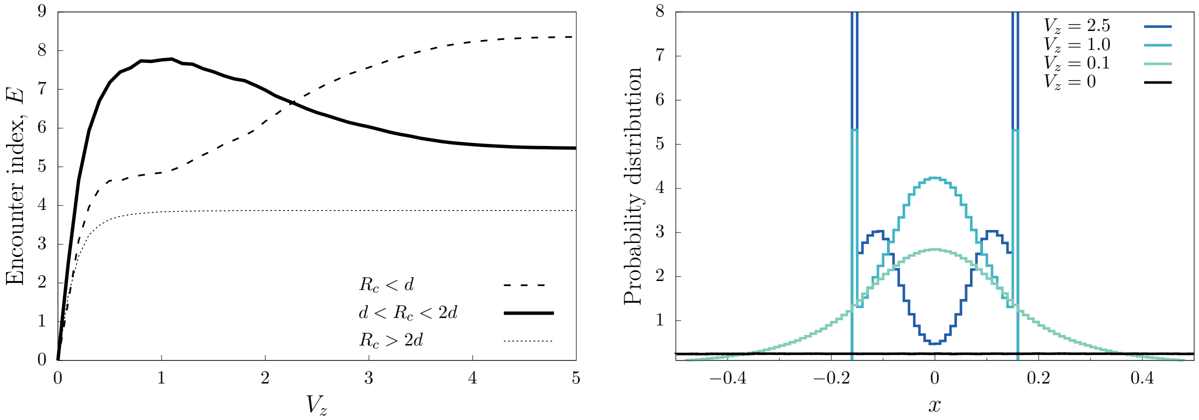


Figure ST2. **Encounter dynamics in the toy model. Left:** encounter index, $E$, as a function of swimming speed. Solid lines represent the encounter index as a function of $V_z$ under low small-scale turbulence ($\sigma_\psi = 0.01$). The different lines correspond to different catching ranges $R_c$ = 0.1 cm, 0.2 cm, 0.4 cm while the distance between prey was fixed at $d = 0.15$. In general, for $R_c \in [d, 2d]$ ($R_c = 0.2$ case), prey availability depends non-monotonically on $V_z$. Therefore, whenever the presence of this configuration is abundant, e.g. low turbulence, a peak in the curve $E(V_z)$ will appear. **Right:** probability distribution for the predator position in the toy model for swimming speeds above and below the optimal one ($V_z \approx 1$). At optimal swimming speed, the predator confines itself in between the two prey, which maximizes the encounter index. Below optimal swimming speed the predator can be found far from the prey individuals, and above the optimal swimming speed the predator is focused only on one of the prey, leading in both cases to values of $E$ that are lower than that corresponding to the optimal speed.

### C. Coarse-graining procedure

In a coarse-graining procedure, a set of steps are followed to scale the interactions occurring at a finer scale to a coarser scale. In our case, we scaled up the interactions observed in the individual-based model (IBM) to obtain a density-field description (DFD) that tracks prey and zooplankton densities $p(r,t)$ and $z(r,t)$ at position $r = (r_x, r_y)$ through time at a coarse spatial scale. To this end, we first divided a large squared system of size $\mathcal{L}$ into pixels of size $L$ compatible with the scale of the IBM. Thus, the coarser system formed a squared-lattice with pixel coordinates given by $r_{m,n} = (x_m, y_n) = (mL, nL)$. For different tuples, $(\sigma_\psi, V_z, L)$, we tracked, for time intervals $\Delta t$, the changes in number of individuals for every pixel, i.e. $\Delta P_{m,n}(t)$ and $Z_{m,n}(t)$. For a given pixel, changes can be due to birth-death events (B) and transport (T), which we represented as $\Delta P_\pm = [\Delta P_\pm^B, \Delta P_\pm^T]$ and $\Delta Z_\pm = [\Delta Z_\pm^B, \Delta Z_\pm^T]$. For the parameters used in our IBM, $\Delta t$ = 1h was a suitable choice that captured the pixel dynamics and was sufficiently short that changes per time step were relatively small (e.g., $\Delta P_\pm^B / P \ll 1$). Therefore, changes within time intervals were negligible when compared to changes across time intervals, and thus variation within time intervals could be neglected.

#### 1. *Demography and interaction terms*

The realized net change for prey abundance can be written as, $\Delta P^B(t) = \Delta P_+^B(t) - \Delta P_-^B(t)$, and zooplankton, $\Delta Z^B(t) = \Delta Z_+^B(t) - \Delta Z_-^B(t)$, with these changes set by the IBM rules. In the case of prey, its spatial distribution does not affect its reproduction, but its mortality stems from feeding and thus the location of zooplankton individuals. Similarly, zooplankton mortality is not dependent on prey, but feeding does depend on the location of prey individuals. Therefore, although the former terms (prey reproduction and zooplankton mortality) can be directly calculated a priori, the latter terms (feeding) depend on emergent properties of the community.

By tracking prey birth and zooplankton mortality events, we first compiled the resulting increase in prey and decrease in zooplankton abundances, $\Delta P_+(t)$ and $\Delta Z_-(t)$ respectively. Then, we represented graphically their corresponding probability distributions; the mean and variance of such distributions are sufficient to characterize their statistical properties. At the scale of the pixel size and for sufficiently large pixels, the probability distribution for changes due to these demographic events can be well approximated by a Gaussian distribution. Finally, we calculated the mean values for such distributions as a function of the corresponding density, which resulted the expected $rp(1-p/K)L^2\Delta t$ term for prey reproduction and a term $dzL^2\Delta t$ for zooplankton mortality (see Fig. ST3), thus matching what would be expected from the rules of our IBM (see parameter definition in main text and *Materials and methods*). These results confirm that our statistical methodology can be used reliably to find the coarser, density-level expressions representing the individual-level interactions. Thus, we followed similar steps to find expressions representing the interspecific interactions, that is, feeding-related terms.

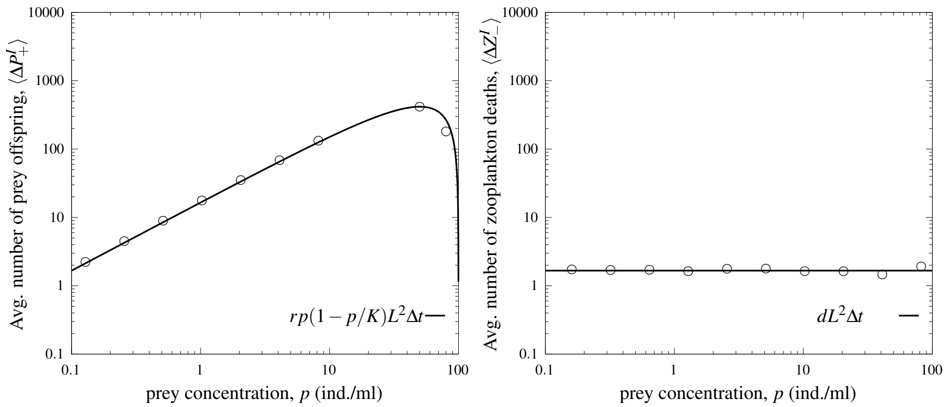


Figure ST3. **Changes in particle number.** Panels show the average offspring number for prey (left) and the average number of zooplankton deaths (right) during $\Delta t = 1$h. Pixel size $L = 20$ cm; other relevant parameter values are listed on Table I. This particular simulation was done assuming moderate small-scale turbulence, $\sigma_\psi = 0.25$ cm/s, and swimming speed $V_z = 0.5$ cm/s, but results qualitatively similar for other values.

We first generalized the expressions for the interaction-related changes in density per unit of time, $F_p \equiv \Delta P^B(t)/(L^2\Delta t)$ and $F_p \equiv \Delta Z^B(t)/(L^2\Delta t)$, by adding a potential feeding term (represented by the function $g(p)$) and stochasticity:

$$F_p(p,z) = rp(1-p/K) - zg(p) + L^{-1}\eta_p\,, \tag{1}$$

$$F_z(p,z) = -dz + bzg(p) + L^{-1}\eta_z\,, \tag{2}$$

The terms $\eta_p$ and $\eta_z$ are Gaussian noises with zero mean and variances given by $\langle\eta_p^2\rangle = |rp(1-p/K)| + zg(p)$ and $\langle\eta_z^2\rangle = bzg(p,z) + dz$. These terms result from the large-number approximation for the birth-death statistics (including feeding), which are implemented as a Poisson process in the IBM. For example, density changes due to feeding in the prey population have the form $\Delta p \sim -\text{Pois}(\mu\Delta t)$, where $\mu = \mu(\{\sigma_\psi, V_z, [p,z], L\}, [\Delta p, \Delta z])\Delta t$ is the mean number of ingestion events within a pixel during time interval $\Delta t$. If pixels are large (e.g., 20 cm as in Fig. 2), the number of events is also large. Then, the Poisson distribution can be approximated by a Gaussian with same mean and variance, $\text{Pois}(\mu\Delta t) \simeq \mathcal{N}(\mu\Delta t, \mu\Delta t)$. Consequently, focusing only on the effects of feeding, the stochastic dynamics of the changes in prey density inside a given pixel can be written as, $\dot{p} = -\mu + L^{-1}\sqrt{\mu}\eta$, where $\eta$ is a zero-mean Gaussian white noise, $\mu = zg(p)$, and $g(p)$ is the feeding function. At large scale, $L \to \infty$, fluctuations can be neglected and thus the dynamics are driven by the average values, leading to the deterministic form of Eqs. 3 from the main text.

Similarly to the steps above for the prey birth and zooplankton mortality, to calculate $g(p)$ from data we collected the values of $\Delta P_-^I$ produced at different densities and environmental conditions, which provided us with a database of the form $\{\{\langle\Delta P_-^I\rangle, \langle(\Delta P_-^I)^2\rangle\}, \{p,z,\mathcal{Q}\}\}$, where $\mathcal{Q} = \{\sigma_\psi, V_z, c\}$ are the IBM parameters characterizing each case. As Figs. S8a-e show, we could also approximate a Gaussian distribution for the feeding terms. The mean in this case was obtained following the observation that, as the system is effectively memory-less due to the timescale separation between relocation and demography (see main text), and feeding depends on the prey individuals available per zooplankton individual as per the IBM rules, the function $g$ should be a function of the encounter index, $E$. Specifically:

$$g(p) = \langle\Delta P_-/Z\rangle = c\pi R_c^2 p(1+E(p)) = g_0(p)(1+E(p)) \tag{3}$$

with

$$E \simeq \frac{\alpha(\sigma_\psi, V_z)}{(p/p_0)^{\beta(\sigma_\psi, V_z)}}\,, \tag{4}$$

Therefore, if $E = 0$, indicating a well-mixed spatial distribution, feeding is reduced to a baseline level $g(p) = g_0(p) = c\pi R_c^2 p$, which is the standard term in the Lotka-Volterra equations. On the other hand, when $E \neq 0$ feeding becomes a function of the environment (in this case, turbulence) but also a function of zooplankton swimming speed $V_z$ and prey density $p$. Therefore, exhaustively measuring the values of $E$ across different settings provides the information needed to parameterize these dependencies and thus the expression above. We observed that $\beta(\sigma_\psi, V_z)$ can be considered roughly constant, $\beta(\sigma_\psi, V_z) \approx 1$, only exhibiting significant changes when $E$ is negligible: high small-scale turbulence and low swimming speed. Contrarily, we found the parameter $\alpha$ to depend more strongly on simulation settings, being well approximated by $\alpha(\sigma_\psi, V_Z) = \alpha_1(\sigma_\psi)(1 - \exp[-V_Z/\alpha_2(\sigma_\psi)])$ with $\alpha_1(\sigma_\psi) = 10.61 \exp(-2.32\sigma_\psi)$ and $\alpha_2(\sigma_\psi) = 6.99\sigma_\psi^{1.66}$ (see Fig. S6).

#### 2. *Transport*

To obtain the transport terms, we followed the same steps as for the interaction terms, now focusing on the impact of adjacent pixels that can send or receive individuals into a focal pixel. Exchange between pixels per unit time can be due to passive or active movement: $\Delta p^T/\Delta t = \Delta p^T_{\text{passive}}/\Delta t$ and $\Delta z^T/\Delta t = \Delta z^T_{\text{passive}}/\Delta t + \Delta z^T_{\text{active}}/\Delta t$. The passive component of transport accounts for the impact of the external flow that affects both species. Small-scale turbulence promotes diffusion with coefficient $D = \sigma_\psi^2/2$, and the large eddies induce advection with a velocity $\vec{v}_f$ that is given by the point-vortex model. These passive contributions to the change due to transport lead to, $\Delta p^T_{\text{passive}}/\Delta t = \partial_t p = L^{-2} D \nabla^2 p - L^{-1} \vec{v}_f \nabla p$ and $\Delta z^T_{\text{passive}}/\Delta t = L^{-2} D \nabla^2 z - L^{-1} \vec{v}_f \nabla z$, where the differential operator is defined as $\nabla p = (p_j - p_i)/\Delta x$ and the lattice spacing $\Delta x = 1$ (since the actual spatial discretization length –or pixel size– $L$ appears explicit in the equations).

The active components can be extracted from the IBM data by measuring the flux, $w_{i,j}$, from pixel $i$ to pixel $j$ due to swimming. Thus, these components are only applicable to zooplankton and only when $V_z \neq 0$. Exchanges occur for individuals that are at the boundaries of the spatial pixel and can experience a gradient of density. For a predator exactly at the edge of an spatial pixel $i$ we can expect that, during a target-selection-and-swimming event, the probability of choosing a target at a neighbor pixel $j$ be approximately $q_{ij} = p_j/(p_i + p_j)$, which can be written as $q_{ij} = [1 + (p_j - p_i)/(p_i + p_j)]/2$. In the limit of small density differences, $q \sim 1 + \nabla p/p$. The constant term accounts for the fact that, even in the absence of any prey density gradient (i.e. $\nabla p = 0$ with $p \neq 0$), swimming would still occur but in a unbiased manner; the second term, which includes $\nabla p/p$, accounts for the environmental cue that triggers swimming, here based on the difference of densities (encoded in the $\nabla$ operator, "chemotactic signal" hereon). This directional term resembles the Keller-Segel model for chemotaxis [4] but with logarithmic sensitivity (Weber–Fechner law): spatial differences are perceived relatively to the local average resource density, an actual ingredient present in our IBM.

Our data (Figs. S8f-h) show that indeed the number of times that swimming originates from a given pixel is given by $qzL^2\Delta t$, where the actual probability, $q = L^{-1}[D_\chi(V_z) + f_\chi(V_z)(\nabla p/p)]$, includes the coefficients $D_\chi(V_z)$ and $f_\chi(V_z)$ representing the weight of each component to the total flux (see Fig. ST4).

Note that both coefficients depend on the swimming speed (and on perceptual range and the frequency of target-selection-and-swimming events, both fixed parameters in our simulations).

The estimate for the probability above leads to the realization that zooplankton movement (i.e. changes in location) can be effectively described by the Langevin equation of a biased random walk [5]: $\dot{\vec{r}} = \vec{v}_\chi + \sqrt{2D_\chi}\vec{\eta}(t)$, where $\vec{v}_\chi = L^{-1} f_\chi \nabla p/p$ ("chemotactic velocity") and $\eta$ is a zero-mean Gaussian white noise with unit variance. From this Langevin equation, we can obtain the associated Fokker-Planck equation [4, 5], whose solution allows us to find the flux of predators $J_z = v_\chi z - D_\chi \nabla z$. The latter therefore provides the last piece of information to calculate the changes of zooplankton density due to movement (i.e. active transport), as $\Delta z^T_{\text{active}}/\Delta t \equiv -L^{-1}\nabla\cdot J_z = L^{-2}\nabla\cdot(D_\chi \nabla z) - L^{-1}\nabla(v_\chi z)$.

Adding together the active and passive components of transport, the changes in density produced by movement for prey and zooplankton, $\Gamma_p$ and $\Gamma_z$, respectively, are given by

$$\begin{aligned} \Gamma_p(p) &= L^{-2} D \nabla^2 p - L^{-1} v_f \nabla p + L^{-1}\xi_p \\ \Gamma_z(p,z) &= \left(L^{-2}D + L^{-2}D_\chi(V_Z)\right)\nabla^2 z - L^{-1} v_f \nabla z - L^{-2} f_\chi(V_Z)\nabla\left(z\left(\nabla p\right)/p\right) + L^{-1}\xi_z\,. \end{aligned} \tag{5}$$

Note that we captured the stochasticity associated with the active movement using zero-mean Gaussian noises, $\langle \xi_p^2 \rangle = L^{-2} D |\nabla^2 p|$ and $\langle \xi_z^2 \rangle = L^{-2} D |\nabla^2 z| + |L^{-2} f_\chi(V_Z) \nabla\cdot(z(\nabla p)/p)|$ (see Fig. S8 and *Materials and methods*).

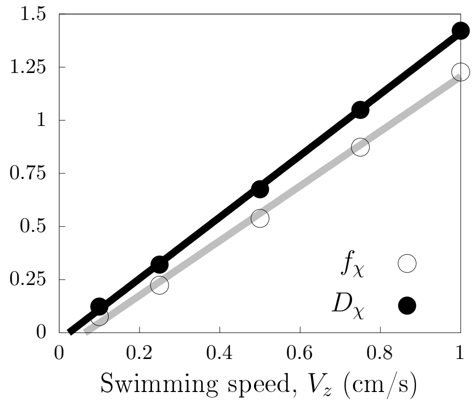


Figure ST4. **Flux between pixels.** Coefficients of the cross-pixel-swimming rate as a function of zooplankton swimming speed. For these plots, we used a 4-by-4 lattice ($L_S = 40$) with pixel size $L = 10$ cm subjected to turbulence intensity $\sigma_\psi = 0.25$ cm/s (eddy structures were neglected for simplicity), fixing densities $(p, z) = (5, 1)$. See Fig. S8 for scaling and probability distribution.

#### *3. Complete density-field description*

Combining the expressions obtained from demographic events and movement, we obtained:

$$\partial_t p(r,t|L,\mathcal{Q}) = F_p(p,z|L,\mathcal{Q}) + \Gamma_p(p|L,\mathcal{Q})\,,$$
$$\partial_t z(r,t|L,\mathcal{Q}) = F_z(p,z|L,\mathcal{Q}) + \Gamma_z(p,z|L,\mathcal{Q})\,, \qquad (6)$$

for a given set of IBM parameters, $\mathcal{Q} \equiv \{\sigma_\psi, V_Z, \ldots\}$, and pixel size $L$. Eq. 6 is thus a DFD for our plankton community that enables studying its dynamics at a wide range of scales by adjusting pixel size $L$.

Note that both transport and birth-death terms depend on the scale of description $L$. For the birth-death events terms, only fluctuations are affected by $L$, whereas the mean component remains unchanged. The contribution of diffusion and "chemotactic" terms, however, decreases as $L$ increases (see Eq. 5). In addition, advection depends on the structure of the velocity field generated by the point-vortex model [6]. The point-vortex model considers several vortices with varied sizes and rotation direction to capture the statistical proprieties of geostrophic turbulence (observed in the 1-100 km range); in this case, velocity intensity exhibits a scaling law, $|\boldsymbol{v}_f| \sim L^{3/2}$ (see Fig. S1 and [7]), and thus advection becomes more important at larger scales, $L^{-1}|\boldsymbol{v}_f| \sim L^{1/2}$ [2].

In summary, in the limit of small pixel size (large $L^{-1}$, i.e. for scales typical of the individual-level interactions), diffusion and "chemotaxis" dominate the dynamics; for very large pixels (which can contain the large eddies), the ecological interaction dominate as "chemotaxis", diffusion, and advection terms become negligible. At scales like the ones corresponding to typical satellite images (~ 1 km), advection and interactions both contribute to large-scale patterns and capture the tug-of-war between physics and behavior that remains represented by nonlinearities in the feeding response function.

### D. Numerical integration of the density-field description

. Eqs. 3 in the main text can be integrated following standard integration schemes. We used a fourth-order Runge-Kutta scheme for the reaction components and a low-order scheme for the transport processes [1], using the free and open source GNU Scientific Library (GSL) for C with a sufficiently small time step (i.e. reducing the time step produces almost identical density values, with a relative error under $10^{-3}$, see documentation in https://www.gnu.org/software/gsl/doc/html/ode-initval.html for details). We used the parallel scheme *openMP* to integrate eddy dynamics. A special remark needs to be made with respect to the integration of the transport terms. When displacing/moving density with a net velocity, $\vec{v}_f$, the new calculated position after a time step will not match exactly a location in the grid due to small numerical differences, regardless of the spatial resolution. To tackle the issue, it is necessary to distribute the displaced density among the closest neighboring pixels. To this end, we followed a backward semi-Lagrangian method where the arriving flow in a given pixel was calculated using a bi-linear interpolation procedure from neighboring pixels [8, 9]. After displacing densities, the total population density can slightly change due to small numerical errors, issue that we fixed by re-scaling density using a factor that keeps total density unchanged [9].

#### E. The 2-species model

In this extension of the main model, we explicitly consider two zooplankton populations, $z_0$ and $z_1$, which represent a passive and an active species, respectively. Prey are now predated upon by both zooplankton populations, which show different feeding rates due to their difference in swimming speed and feeding behavior. Thus, under an upwelling scenario like the one described in the main text, the model equations are:

$$\begin{aligned}
\partial_t p(r,t) &= rp(1 - p/K) - (z_0 g_0(p) + z_1 g_1(p)) + L^{-1} v_f \nabla p\,, \\
\partial_t z_0(r,t) &= -d_0 z + b z_0 g_0(p) + L^{-1} v_f \nabla z\,, \\
\partial_t z_1(r,t) &= -d_1 z + b z_1 g_1(p) + L^{-1} v_f \nabla z\,, \\
\partial_t K(r,t) &= -\tau^{-1}(K - \bar{K}) + L^{-1} v_f \nabla K\,,
\end{aligned} \tag{7}$$

An expected trade-off between behavioral strategies can be implemented by modifying the parameters of the equations. For the results shown in Fig. S13, for simplicity we assumed that both zooplankton are paying an identical cost ($d_0 = d_1$) but due to different behaviors: while the active predator follows an informed feeding strategy, $g_1 = c\pi R_c^2 p(1 + E(p))$ (with $V_z = 0.5$), the passive predator is non-motile and constantly scanning its surroundings, $g_0 = c_0 c\pi R_c^2 p$ (we set $c_0 = 10$ in Fig. S13). Other parameters are discussed in the main text and specified in the Fig. S13 caption. The figure shows the 2-species version of Fig. 4, for which the adaptive-feeding model was used instead. The results of the 2-species model and the adaptive feeding model are qualitatively similar, with sharpness increasing for low prey density. Quantitatively, however, note that since behavior dominance in the 2-species model requires one species to outgrow the other, there is a delay between changes in local prey density and changes in the degree of dominance of the active strategy (quantified as $100\cdot(z_1 - z_0)/z_0$); this delay, together with the continuous mixing process, enables local mismatches between dominance and local prey densities (see Fig. S13), something that does not happen in the adaptive-feeding model. As another consequence, the threshold prey density for which there is an abrupt change in zooplankton behavior differs for both models, with shifts towards lower densities for the 2-species cases.

[9] G. P. Brasseur and D. J. Jacob, *Modeling of atmospheric chemistry* (Cambridge University Press, 2017).

| | | |
|---|---|---|
| $r$ | prey reproduction rate | 1.0 (day$^{-1}$) |
| $d$ | zooplankton death rate | 0.1 (day$^{-1}$) |
| $c$ | zooplankton search rate | 10 – 100 (individuals$^{-1}$day$^{-1}$) |
| $b$ | zooplankton energy conversion factor | 0.01 (prey/zooplankton individuals) |
| $K$ | prey carrying capacity | 100 – 10000 (cm$^{-2}$) |
| $R_c$ | zooplankton catching range | 0.2 (cm) |
| $R_p$ | zooplankton perceptual range | 0.5 (cm) |
| $V_z$ | zooplankton swimming speed | 0 – 1.0 (cm/$s$) |
| $\sigma_\psi$ | small-scale turbulence intensity (speed std. dev.) | 0 – 1.0 (cm/$\sqrt{s}$) |
| $\psi$ | eddies maximum angular speed | 0 – 1.0 (day$^{-1}$) |
| $p$ | prey density | Variable (individuals cm$^{-2}$) |
| $z$ | zooplankton density | Variable (individuals cm$^{-2}$) |
| $P$ | number of prey individuals | Variable (individuals) |
| $Z$ | number of zooplankton individuals | Variable (individuals) |

Table I. List of parameters and variables used in the IBM. Parameter values were chosen to resemble known prey and the zooplankton that predate on them (e.g. dinoflagellates, diatoms, and microzooplankton for the former, copepods and fish larva for the latter), considering the size of the zooplankton to be around 0.1cm. For more details see *Materials and methods*—Individual-based model description and parameterization.

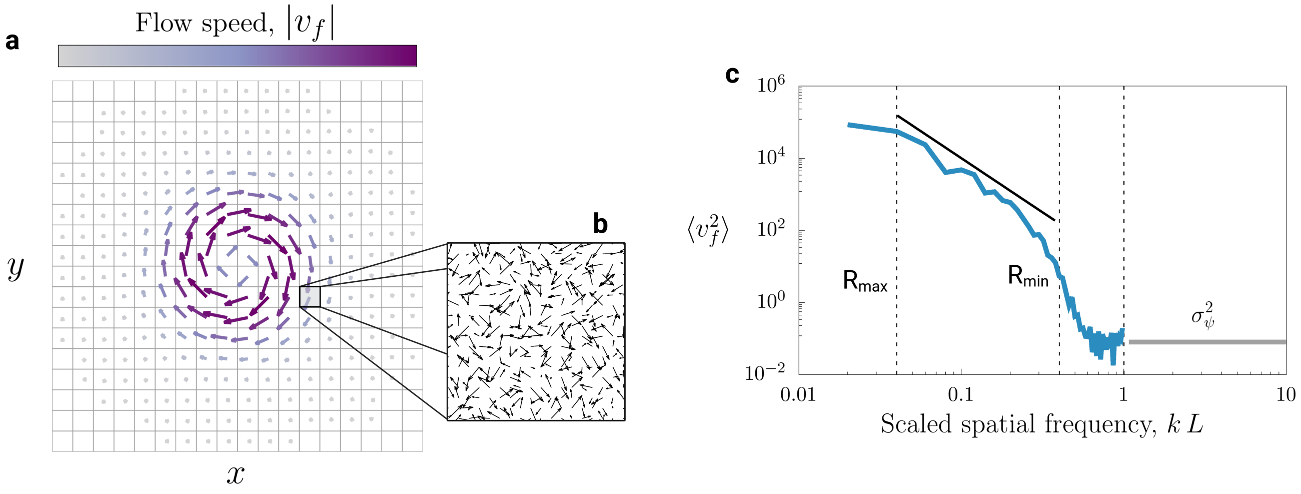


Figure S1. **Turbulence model.** In our multiscale framework, we used two models to describe the flow velocity field at small and large scales. At large scales, we used a point-vortex model that accounts for many eddies with different size and rotation direction to generate the realistic features of turbulence. In panel a) the vector field show the velocities induced by a single eddy. Colors indicate the intensity of the speed, which reaches a maximum for intermediate distance from the eddy center. For spatial scales below the pixel size, we used a simplified flow model where the velocities are random vectors (see *Materials and methods*), as shown in panel b. The combination of both models allow for the continuity of the flow velocity spectrum as shown in panel c). For large scales, the velocity field is structured and characterized by a power-law scaling (see blue line produced using the point-vortex model) that is typically observed in the ocean [7]. At small scales, any coherent structure is expected to be fade leaving tiny velocity fluctuations, which are preserved by the random flow model (gray solid line). Vertical dashed lines highlight eddy maximum and minimum sizes, $R_{\max}$ and $R_{\min}$ respectively, and $L$, the pixel size (which is 1 in the scaled horizontal axis in panel c). The above results use turbulence intensity $\psi = 0.6$, leading to an average speed of 1km/day in a 120m square domain and $\sigma_\psi = 0.25$ cm/s.

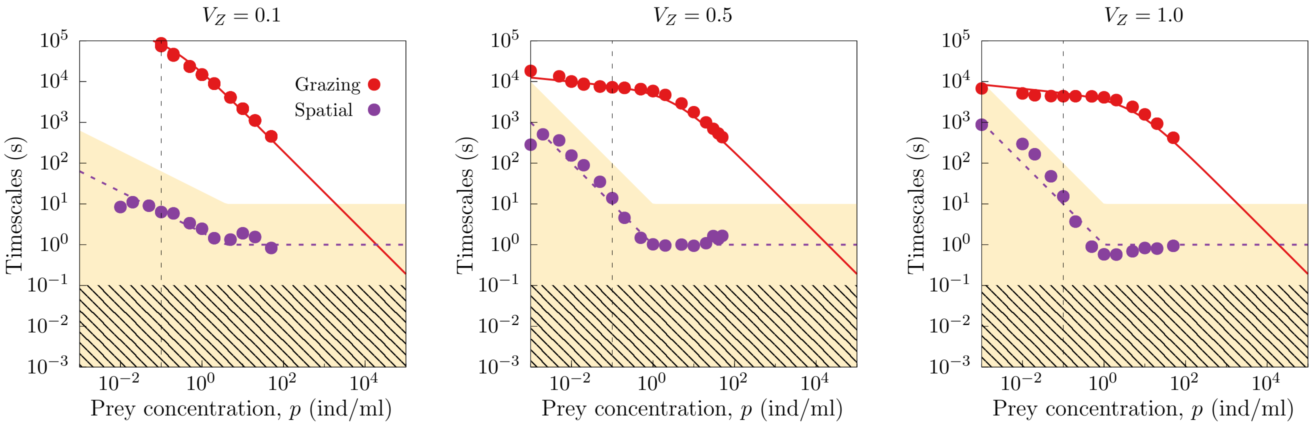


Figure S2. **Timescale separation analysis.** Red and purple dots show the values of the feeding and spatial timescales, in seconds, for $V_z = 0.1$ (left panel), $V_z = 0.5$ (middle panel), and $V_z = 1.0$ cm/s (right panel). Feeding timescale is defined as the average inter-ingestion time, and spatial timescale is defined as the characteristic relaxation times of the spatial index $E$ (see Fig. S4). We can conservatively define the condition for timescale separation as spatial relaxation being 10 times faster than feeding, i.e. when feeding dots are inside the white region in the figure. In this regime, which occurs approximately for $p \in [10^{-3}, 10^{4}]$ in our $V_z = 0.5$ reference case, we can approximate the encounter index as $E(t) \simeq E(p)$. Results shown are for small-scale turbulence used in the manuscript figures, $\sigma_\psi = 0.25$ cm/s. Lines show the theoretical expectations for the behavior of the timescales as a function of prey density: red solid line (feeding timescale) is directly given by $1/g(p)$ while the purple dashed line represent the scaling relation, $1/p^{\gamma}$, where we selected $\gamma = 0.5$ for $V_z = 0.1$ and $\gamma = 1.0$ for $V_z = 0.5$ and 1.0, to guide the trend and provide a visual boundary to the region with timescale separation. System size was chosen within the range from $L = 20$ to $L = 500$ cm to ensure the presence of at least a few hundreds of prey individuals. As shown in Fig. ST1, this timescale separation is *not* an artifact of the simulation algorithm, that is, it does not result from an unrealistic "waiting without eating" period between demographic events. Note that, for the passive zooplankton case $V_z = 0$, there is no spatial structure and therefore the spatial timescale is zero.

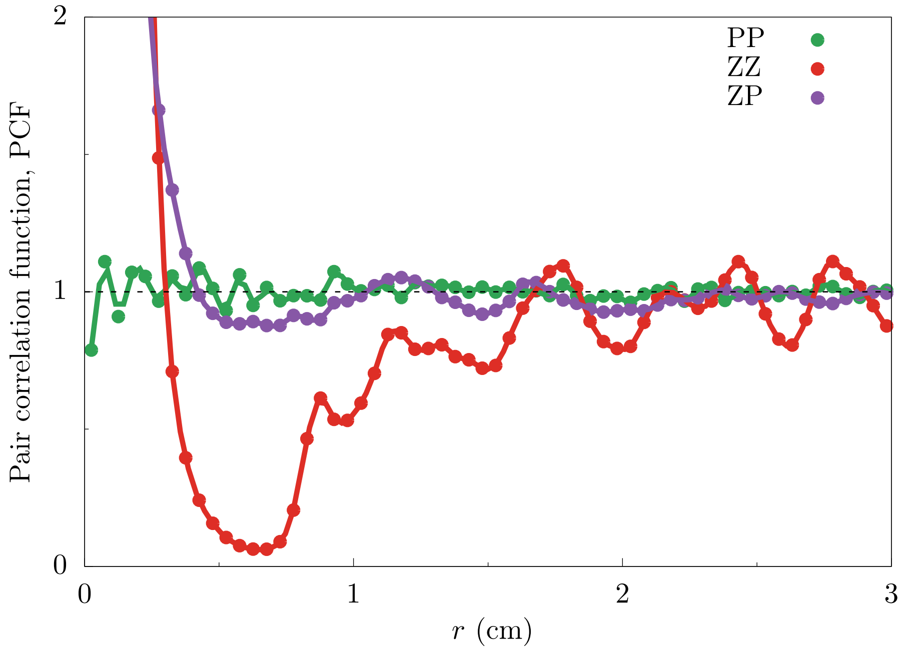


Figure S3. **Pair-correlation function.** Pair-correlation function (PCF) between prey (PP), zooplankton (ZZ), and between zooplankton and prey (ZP) for the case shown in Fig. 2a. where $(\sigma_\psi, V_z) = (0.25, 0.5)$ and $L = 20$ cm (see Fig. 2a caption for more details). For both ZZ and ZP, short scales lead to a substantial positive deviation from the well-mixed scenario (PCF = 1) that indicates: i) higher interspecific encounter rate and ii) zooplankton aggregation. The spatial correlation given by the PP curve, on the other hand, indicates that prey are fairly well-mixed (PCF $\simeq$ 1).

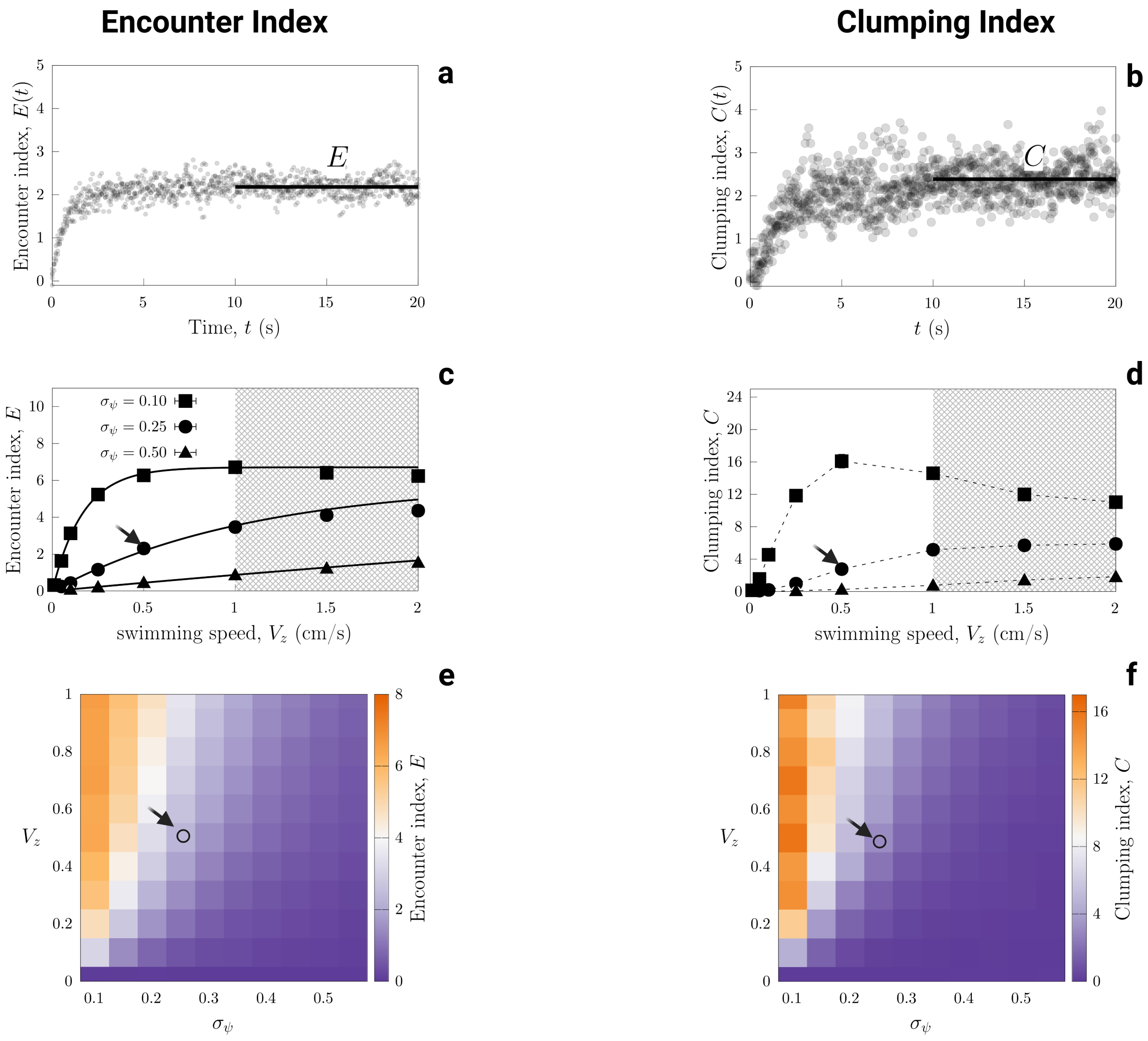


Figure S4. **Encounter and clumping index transient and stationary values.** Panels show the transient (a-b) and stationary values (c-f) for the encounter index $E(t)$ (left column) and clumping index $C(t)$ (right column), for different swimming speeds and small-scale turbulence intensity, $\sigma_\psi$. In the first row (a-b), for $(\sigma_\psi, V_z) = (0.25, 0.50)$, the stationary values $E$ and $C$ for the spatial indices are highlighted with a solid black line, values that are achieved at characteristic time scales 0.8 and 1.8 seconds, respectively. Second and third rows summarize the dependence of $E$ (c &e) and $C$ (d &f) on swimming speed, $V_z$, and small-scale turbulence intensity, $\sigma_\psi$. The heatmaps e-f illustrate the tug-of-war between $V_z$ and $\sigma_\psi$ on the emergence of correlations among individuals. The specific case $(\sigma_\psi, V_z) = (0.25, 0.50)$ shown in a-b is indicated with an arrow in other plots. Simulations were performed starting from the well-mixed state ($E = 0$) with fixed densities, $p = 1.28$ and $z = 1$, as in Fig. 2a-b (see Fig. 2 caption for remaining parameter values).

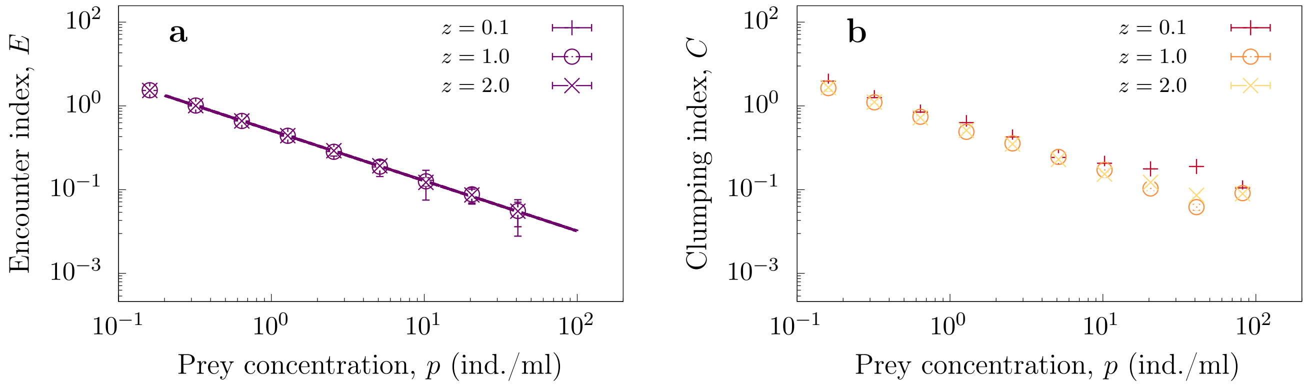


Figure S5. **Encounter and Clumping index as a function of** $(p, z)$**.** Panels a-b complement Fig. 2c showing both spatial indices as a function of prey density for different zooplankton densities, for a fixed swimming speed $V_z = 0.5$ cm/s. Results for both indices remain robust against changes in zooplankton density $z$, leading to the conclusion that only the dependence on $p$ is of relevance. Moreover, the solid line in (a) shows that the power-law of Eq. 1 in main text is valid regardless of zooplankton density $z$. See Fig. 2 caption for details.

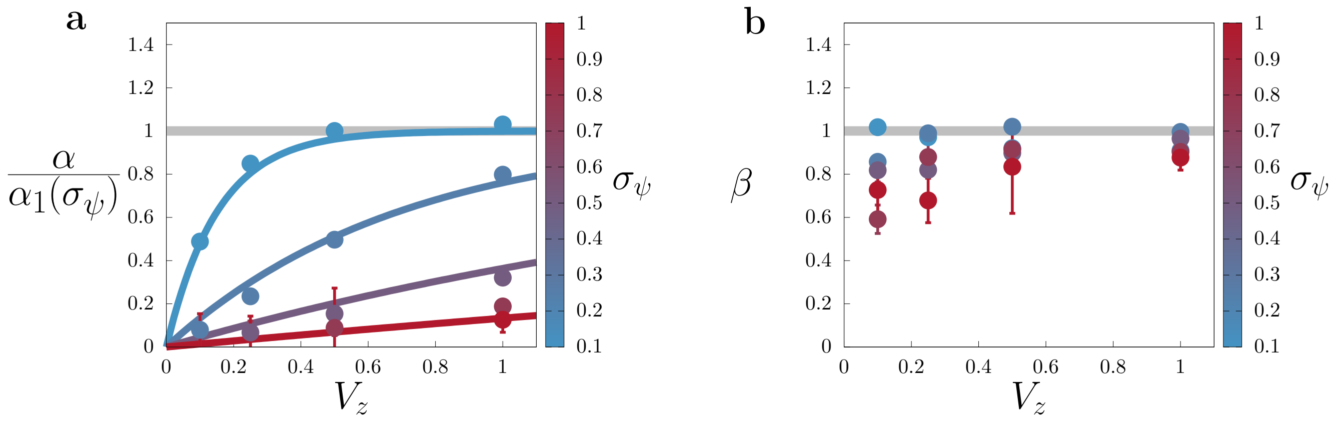


Figure S6. **Dependence of the** $\alpha$ **and** $\beta$ **functions on** $V_z$ **and** $\sigma_\psi$**.** a) Parameter $\alpha(\sigma_\psi, V_Z) = \alpha_1(\sigma_\psi)(1 - \exp[-V_Z/\alpha_2(\sigma_\psi)])$ (scaled by its amplitude, $\alpha_1$) for different turbulence intensities $\sigma_\psi$ (color bar), with $\alpha_1(\sigma_\psi) = 10.61 \exp(-2.32\sigma_\psi)$ and $\alpha_2(\sigma_\psi) = 6.99\sigma_\psi^{1.66}$. b) Exponent $\beta$ as a function of swimming speed $V_z$, for different turbulence intensities $\sigma_\psi$ (color bar).

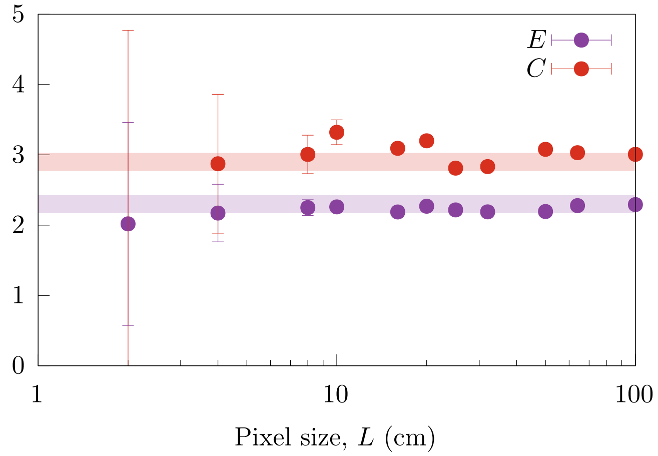


Figure S7. **Spatial indices become invariant beyond $L \simeq 10$ cm.** Encounter and clumping indices as a function of pixel size $L$ (dots). Beyond $L \simeq 10$, the standard deviations (error bars) vanish and the values of the spatial indices achieve the convergence values, indicated by thick solid lines. For this plot, $(p, z) = (1, 5)$ and $(\sigma_\psi, V_z) = (0.25, 0.5)$, representing one of the cases in Fig. 2b (where $L = 20$ was used); see Fig. 2 caption for rest of simulation parameters.

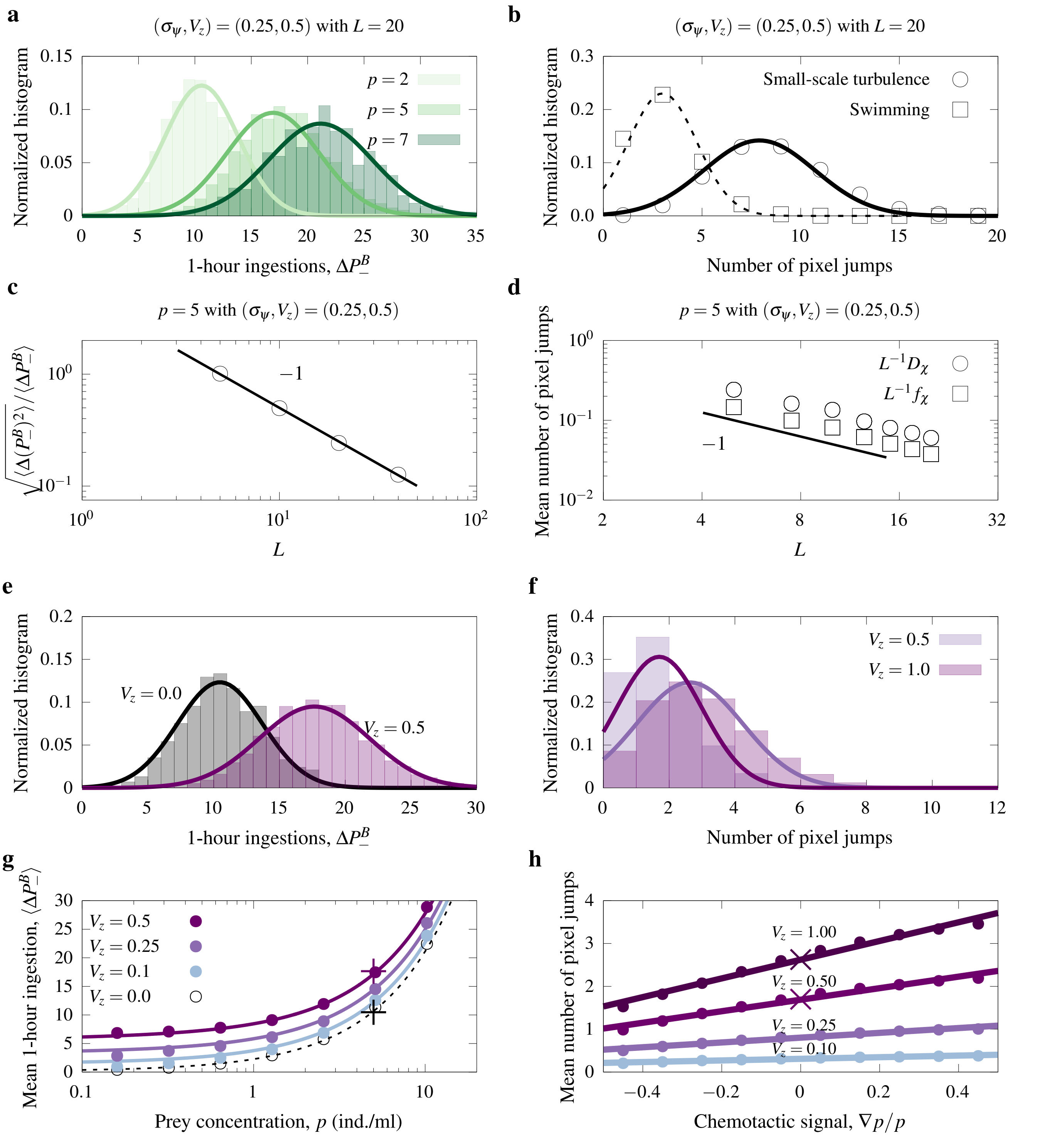


Figure S8. **Coarse-scale statistics.** (a-b) Normalized histogram for the number of prey ingested during 1 hour, under different (fixed) prey densities, within a pixel size $L = 20$ cm (panel a); and for the number of cross-pixel-swimming events due to small-scale turbulence and swimming (panel b). (c-d) Size-scaling for the coefficient of variation for ingestion (panel c) and the mean number of cross-pixel-swimming events (panel d) for $p = 5$ ind/cm$^2$. (e-f) Normalized histogram for the number of prey ingested during 1 hour (panel e) and for the number of cross-pixel-swimming events due to swimming (panel f) for $V_z = 0.0$ cm/s (black) and $V_z = 0.5$ cm/s (purple) and homogeneous environments (thus $\nabla p \approx 0$). g) Mean 1-hour ingestion for the case of panel (e) as a function of prey density, $p$, for different swimming speeds, $V_z$. Crosses mark the case represented in panel e. h) Mean number cross-pixel-swimming events as a function of "chemotactic signal", $\nabla p / p$ for different swimming speeds. Crosses mark the $\nabla p / p \approx 0$ case of panel f. Solid and dashed lines in a,b,e,f are Gaussian fits assuming equal mean and variance (an approximation for the Poisson distribution). Small-scale turbulence intensity is $\sigma_\psi = 0.25$ cm/s, search rate $c = 1.0$ day$^{-1}$, $L = 20$ cm, and species density $(p, z) = (5, 1)$, unless otherwise specified. See *Supplementary Text* for details.

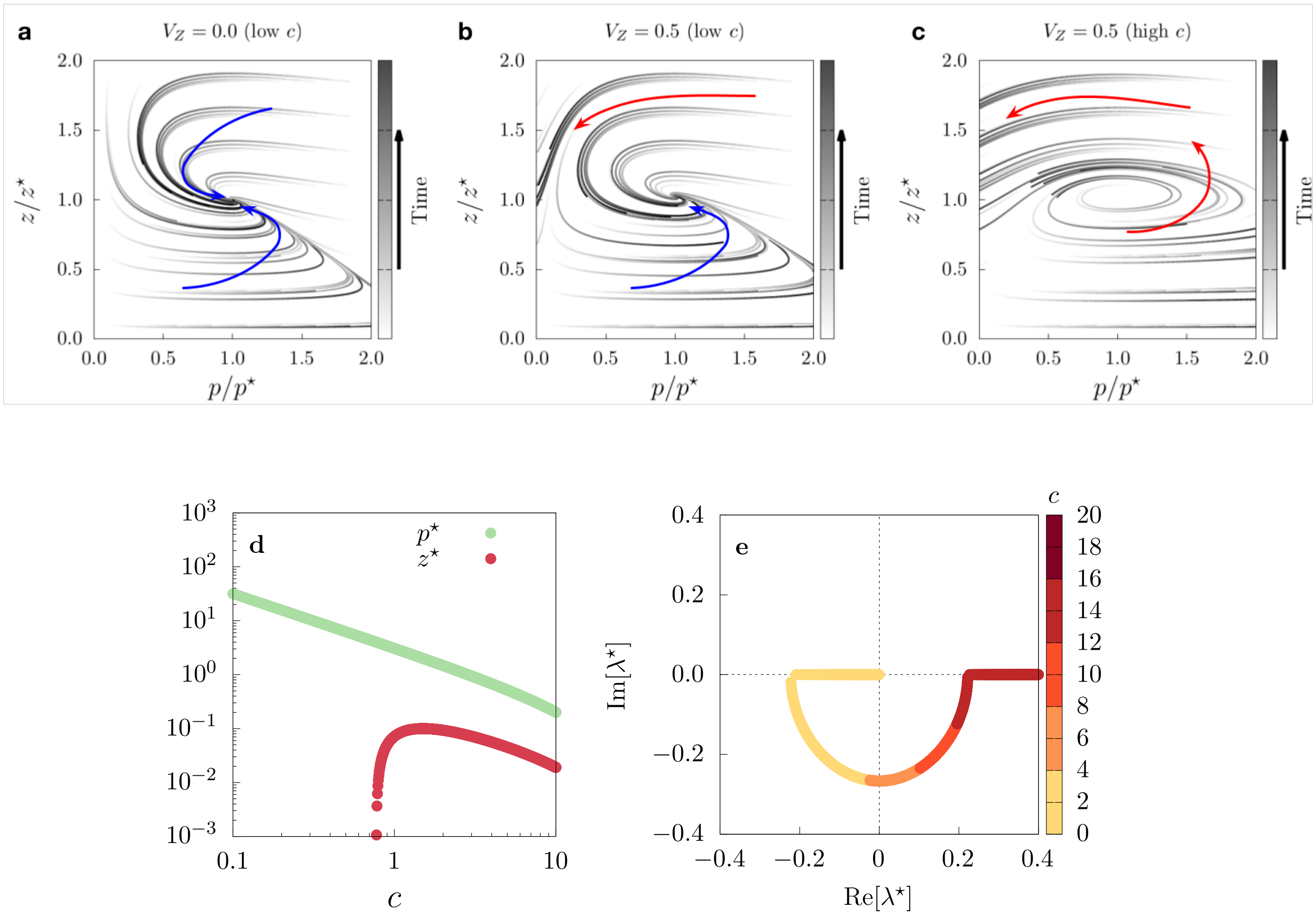


Figure S9. **Active behavior leads to unstable community dynamics.** Trajectories for the passive ($V_z = 0.0$, panel a) and active zooplankton ($V_z = 0.5$ cm/s, panels b and c) cases with low search rate $c = 2$ day$^{-1}$ (panels a-b) and high search rate, $c = 5$ day$^{-1}$ (panel c). Trajectories were obtained numerically integrating, $\dot{p} = F_p(p, z)$ and $\dot{z} = F_z(p, z)$ (Eqs. 1 and 2 without the noises), with different initial conditions. Time increases from white to black color, with arrows highlighting the trajectories that lead to coexistence (blue) and extinction of one of the species (red). Results show that, as active behavior becomes more relevant (i.e. from panel a to c), the community dynamics become more unstable and thereby prone to extinction. This conclusion is supported by the linear stability analysis shown in panels d-e as follows. d) The system has only one (non-trivial) homogeneous state $(p^\star, z^\star)$, shown as a function of the predator search rate, $c$. e) Performing a small perturbation around the homogeneous state, we obtained the real and imaginary parts of the dominant eigenvalue (eigenvalue with the largest real part); positive and negative real part suggest outward and inward spirals, respectively, with the value of the imaginary part corresponding to the angular velocity around $(p^\star, z^\star)$. This linear stability analysis allowed us to predict the qualitative dynamics shown in panels a-b-c.

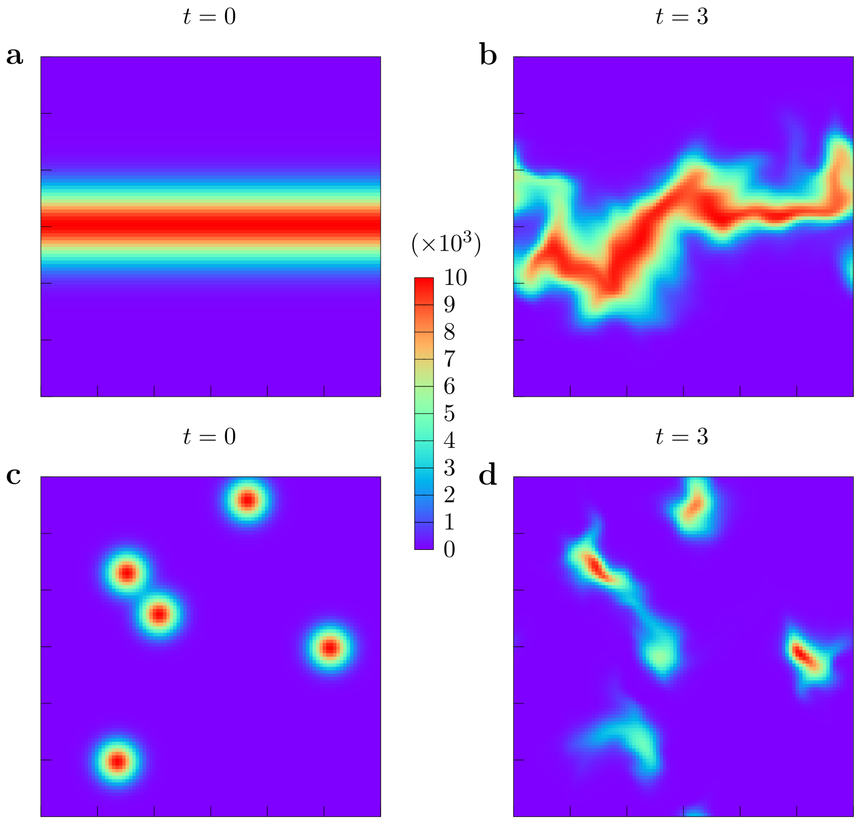


Figure S10. **Different ways to simulate upwelling events.** Carrying-capacity profiles, $K(r,t)$ for the band (top row) and randomly distributed hotpots of nutrient input (bottom row) scenarios at $t = 0$ (left column) and at $t = 3$ days (right column). Colors indicate the carrying capacity value. The steps to simulate the nutrient band profile are described in the main text; for the random hotspots, we simulated the input of nutrient at random locations by using a Gaussian profile for the carrying capacity (i.e. carrying capacity decreasing, from a maximum of $10^4$, following a Gaussian exponential function of the distance to the hotspot center with variance $\sigma^2 = 15$).

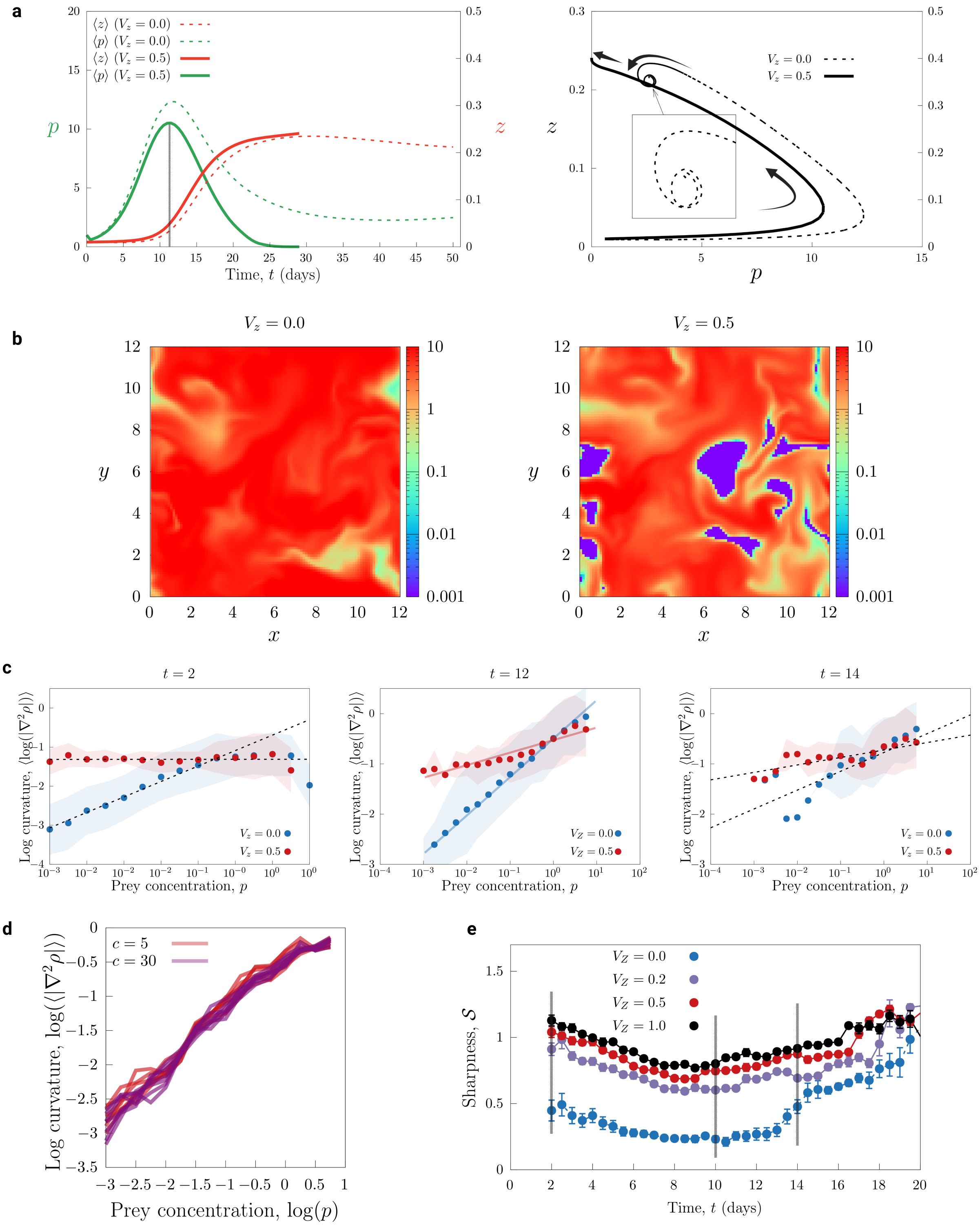


Figure S11. **Dynamics of sharpness.** a) Spatial average of prey density $\langle p \rangle$ and zooplankton density $\langle z \rangle$, as a function of time (left panel) and corresponding trajectory in the $(p, z)$-space (right panel). In both panels, solid lines represent results for $V_z = 0.0$ and dashed lines results for $V_z = 0.5$ cm/s. Initial condition was $p(x, y, t = 0) = 1 \ll K = 100$ and $z(x, y, t = 0) = 0.01$ ind/cm$^2$. b) prey density at $t = 12$d (as marked in a) for $V_z = 0.5$. Color indicates density, in ind/mL, in log-scale to highlight the differences when predators are active. See Fig. 3 caption for simulation details. c) Curvature as a function of prey density for $V_z = 0$ and $V_z = 0.5$ for three snapshots at $t = 2$, $t = 12$ and $t = 14$ d (from left to right). The three snapshots show a wide range of densities that led to a clear power-law scaling, still observed at $t = 14$ d. For very long times, for which the upwelling event is in a fading phase, deviations from a power law are expected. Snapshot $t = 12$ corresponds to the peak of prey density during the upwelling event, for which prey density spatial distribution is shown in log-scale in Fig. 3. d) Curvature-density relation considering a passive predator, $V_z = 0.0$, obtained for different values of the search rate $c$ (colors) and different days around the peak of the upwelling event within the observation window (different lines, $t$ = 12-14 d). e) Prey pattern sharpness as a function of time for simulation data shown in panel c, with error bars representing standard deviation for sharpness when calculating it from curvature data.

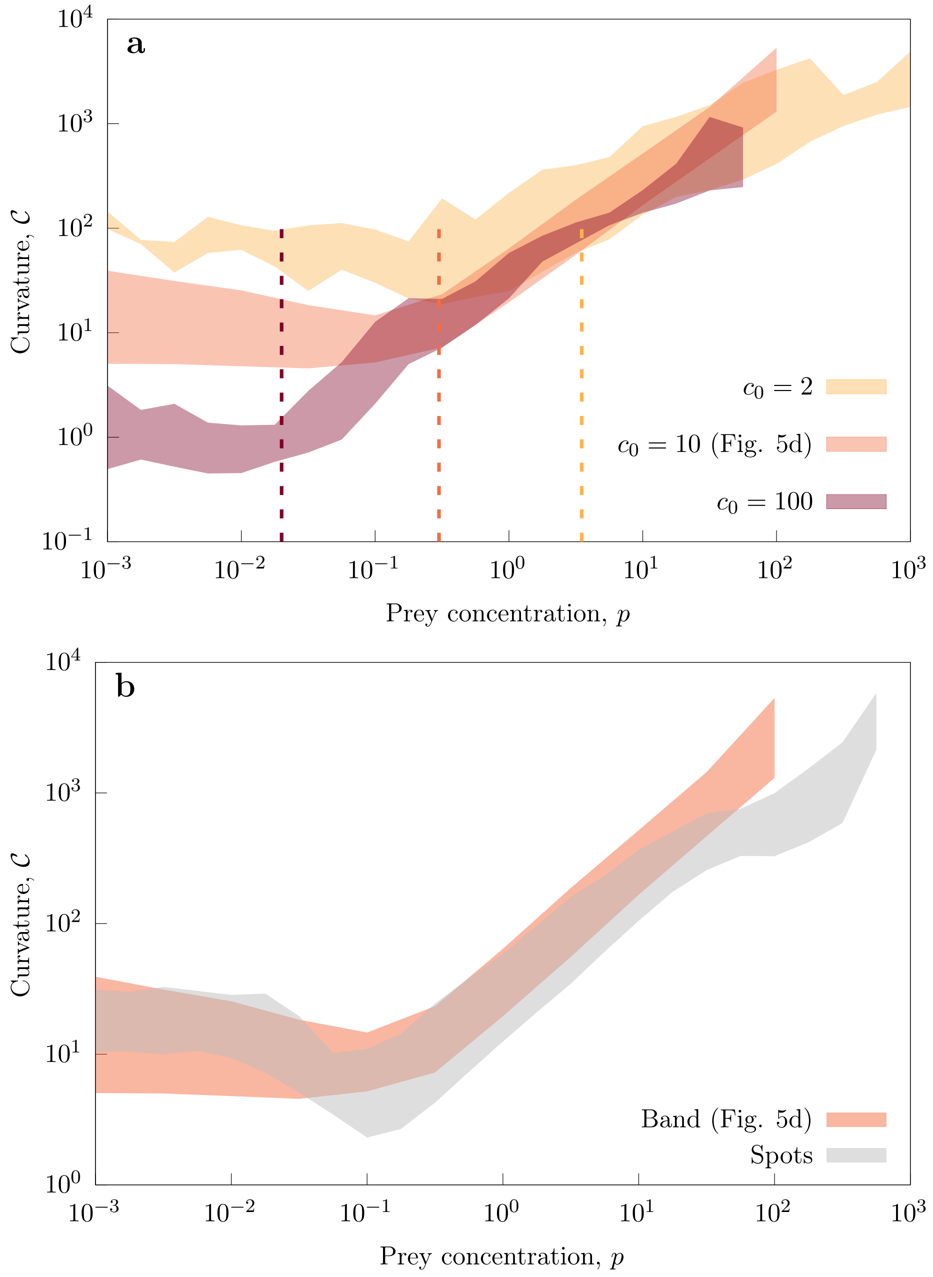


Figure S12. **Curvature-density relation sensitivity analysis.** a) Curvature-density relation for passive-zooplankton parameter $c_0$, used in the adaptive-feeding model. The vertical dashed lines show the density threshold below which feeding actively is more beneficial than feeding passively, which these data show decreases as $c_0$ increases. b) Curvature-density relation for both the nutrient band case (in red, as in a) and Fig. 4d) and the random hotspot case (in gray).

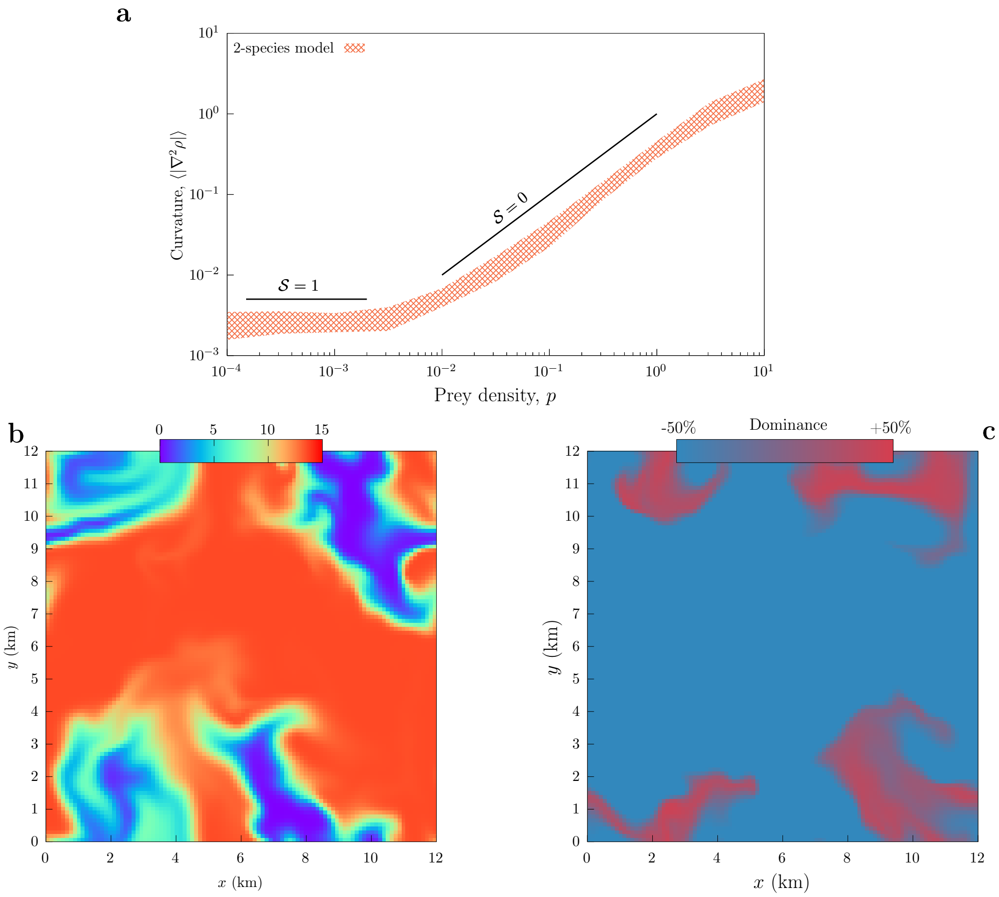


Figure S13. **Spatial niche partitioning (2-species model).** a) Curvature-density relationship extracted from the 2-species model. Solid black lines ($\mathcal{S}$ = 0 and $\mathcal{S}$ = 1) indicate the slope, associated with dominance of a particular feeding behavior. b) Prey density and c) dominant-zooplankton-behavior patterns for the 2-species model with $V_z$ = 0.5 cm/s, $q$ = 4, $d_1$ = $d_0$ = 0.10 day$^{-1}$, $c_0$ = 10 (chosen using ecological arguments, see *Materials and methods* for details). Remaining parameters are the same as in Fig. 3a. Dominance is defined as the relative difference between active and passive zooplankton densities, $100 \cdot (z_1 - z_0)/z_0$. Thus, +50% dominance implies 50% more abundant active predators than passive at that specific time and location.
